# A unified model for cell-type resolution genomics from heterogeneous omics data

**DOI:** 10.1101/2024.01.27.577588

**Authors:** Zeyuan Johnson Chen, Elior Rahmani, Eran Halperin

## Abstract

The vast majority of population-scale genomic datasets collected to date consist of “bulk” samples obtained from heterogeneous tissues, reflecting mixtures of different cell types. In order to facilitate discovery at the cell-type level, there is a pressing need for computational deconvolution methods capable of leveraging the multitude of underutilized bulk profiles already collected across various organisms, tissues, and conditions. Here, we introduce Unico, a unified cross-omics method designed to deconvolve standard 2-dimensional bulk matrices of samples by features into 3-dimensional tensors representing samples by features by cell types. Unico stands out as the first principled model-based deconvolution method that is theoretically justified for any heterogeneous genomic data. Through the deconvolution of bulk gene expression and DNA methylation datasets, we demonstrate that the transferability of Unico across different data modalities translates into superior performance compared to existing approaches. This advancement enhances our capability to conduct powerful large-scale genomic studies at cell-type resolution without the need for cell sorting or single-cell biology. An R implementation of Unico is available on CRAN.

## 1 Introduction

Studying cell-type level genomic variation is critical for unveiling complex biology. Unfortunately, collecting large and well-powered datasets at cell-type resolution for population studies has yet to become common practice. Current single-cell datasets typically consist of data collected from no more than several dozens of individuals due to prohibitive costs, and purifying cell types at scale using flow cytometry is laborious and often impractical, particularly for solid and frozen tissues for which cell isolation is very challenging [1–5].

Indeed, most transcriptomic and other genomic data types collected to date have been measured from heterogeneous tissues that consist of multiple cell types, resulting in vast amounts of large heterogeneous “bulk” genomic data (e.g., over two million bulk profiles publicly available on the Gene Expression Omnibus alone [6]). This rationalizes the development of computational methods that can disentangle the convolution of cell-type level signals that compose such bulk profiles. The premise, upon successful implementation, offers a transformative capability to conduct powerful, large-scale studies at the cell-type level in multiple tissues and under numerous conditions for which large bulk data have already been collected.

Here, we propose a method for deconvolving 2-dimensional (2D) bulk data (samples by features) into its underlying 3-dimensional (3D) tensor (samples by features by cell types). Thus far, deconvolution methods have been tailored to specific data types [7–11]. In contrast, we introduce a unified cross-omics method, Unico, the first principled model-based deconvolution method that is theoretically applicable to any heterogeneous genomic data. As we demonstrate through a comprehensive analysis of multiple gene expression and DNA methylation datasets, this generalization translates into superior performance over existing approaches and improves our ability to conduct powerful large-scale genomic studies at cell-type resolution.

## 2 Results

### From bulk genomics to cell-type resolution: decomposition versus deconvolution

The study of bulk genomics routinely calls for *decomposition*, wherein an observed bulk data matrix is modeled as the product of two matrices: (i) cell-type proportions (fractions) of the samples in the data and (ii) per-feature cell-type genomic levels (“signatures”; Figure 1a). This amounts to solving a matrix factorization problem. For a given bulk observation *x_ij_* of genomic feature *j* in sample *i*, virtually all decomposition models share the following assumption:

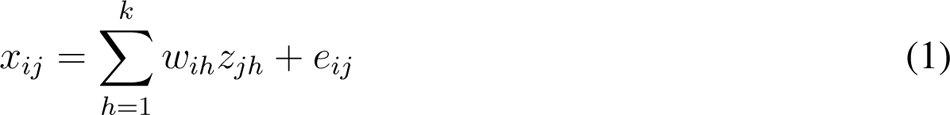

where *w_i_*_1_*, …, w_ik_* are the proportions of *k* modeled cell types in sample *i*, *z_j_*_1_*, …, z_jk_* are the cell-type level signatures of the genomic feature *j* in each of the *k* cell types, and *e_ij_* is an error term.

**Figure 1:**
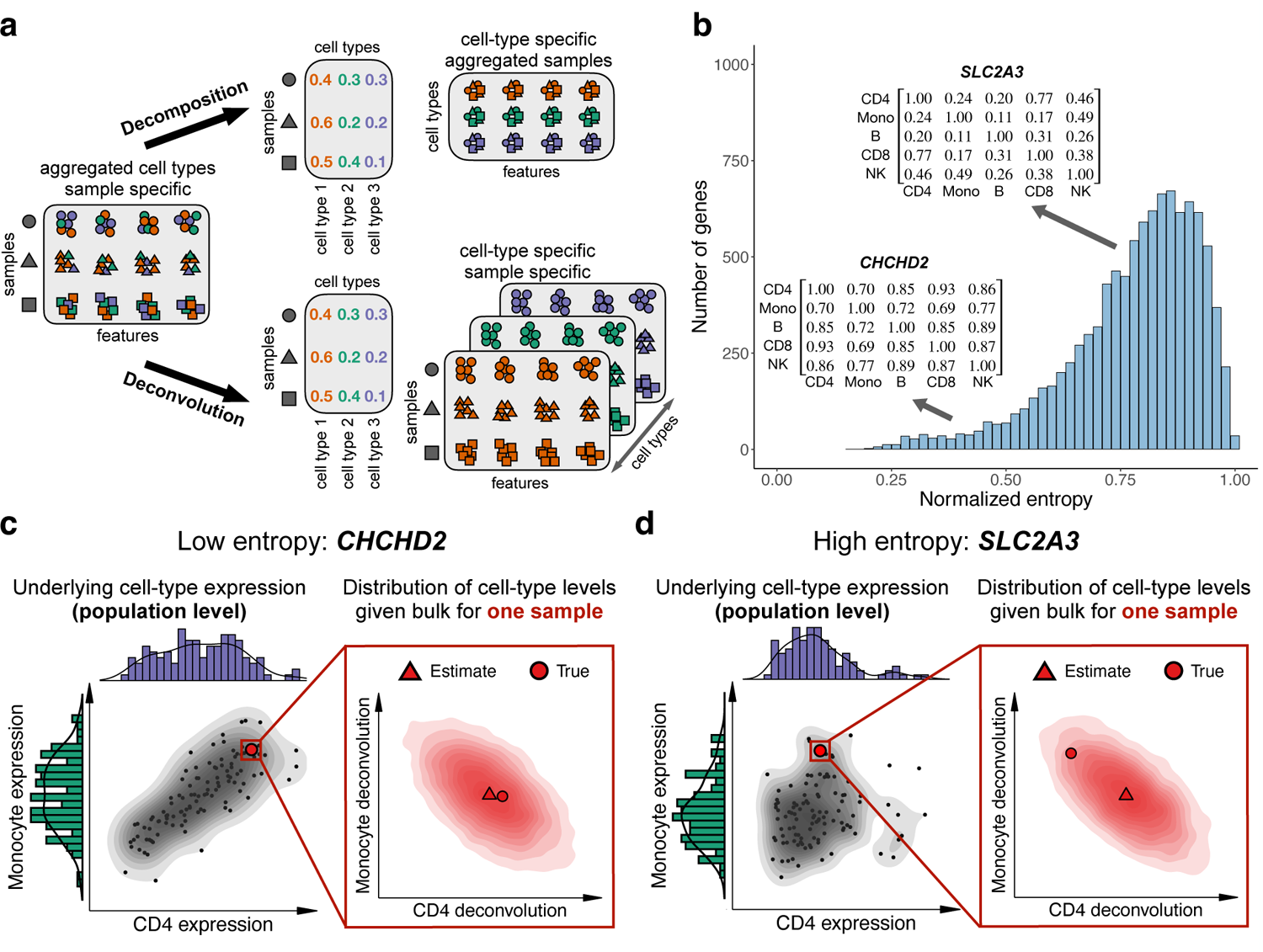
(a) Illustration of decomposition versus deconvolution. (b) Cell-type covariance strength across the top 10,000 most highly expressed genes in scRNAseq from PBMC [29], measured by normalized von Neumann entropy (Methods). (c) The joint distribution of CD4 and monocyte expression in the low-entropy gene CHCHD2 across 118 scRNAseq PBMC samples [29] (left). For one arbitrary sample, Unico’s estimated distribution of the possible CD4 and monocyte expression levels of the sample provides an accurate point estimate based on the sample’s CHCHD2 bulk expression (right). (d) Same as (c), only for the high-entropy gene SLC2A3, providing less cell-type covariance information for the deconvolution.

Numerous decomposition formulations with various assumptions on the products of the factorization have been proposed for the estimation of cell-type compositions and for learning cell-type signatures using different genomic modalities, including gene expression [12–15], DNA methylation [16–20], copy number aberrations [21, 22], ATAC-Seq [23], and Hi-C data [24]. The rich toolbox of decomposition methods has proven successful for a wide range of applications, such as clustering genes and studying their functional relationships [25, 26], inferring tumor composition [21, 22], and discovering cancer sub-types [27]. However, these methods allow us to infer only a single profile of cell-type level signatures per feature, which corresponds to the unrealistic assumption that all samples in the data share the same genomic levels at the cell-type level [28].

Every sample may reflect its own – possibly unique – cell-type level patterns, owing to various factors of inter-individual variation, such as genetic background, environmental exposures, and demographics. A natural adjustment of the decomposition model to reflect such variation yields:

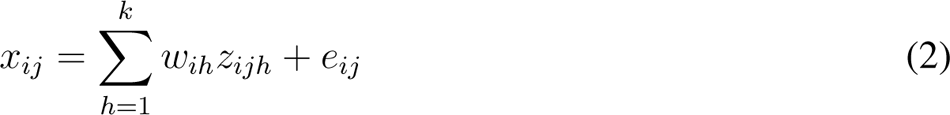

where *z_ijh_* now represents the level of feature *j* in cell-type *h*, specifically in sample *i*. Learning *z_ijh_* from bulk data is essentially a *deconvolution* problem, wherein we disentangle the mixture of signals in a 2D samples by features bulk data into the unobserved underlying 3D tensor of samples by features by cell types (Figure 1a).

Decomposition under Equation (1) can be viewed as a degenerate case of the more general deconvolution problem in Equation (2) [28]. *Deconvolving* the data is thus more desired than merely *decomposing* the data, and the higher resolution of a successful deconvolution is expected to improve cell-type context and discovery in the analysis of bulk genomics. This has been highlighted and demonstrated by several recent deconvolution methods, including CIBERSORTx [8], MIND [9], bMIND [10], and CODEFACS [11] in the context of transcriptomics and TCA [7] in the context of DNA methylation.

### Unico: A unified cross-omics deconvolution model

Current deconvolution methods can be categorized into two groups: heuristic approaches, including CIBERSORTx [8] and CODEFACS [11], and methods based on the assumption of data following a normal distribution, including TCA [7], MIND [9], and bMIND [10]. The latter group faces limitations rising from the normal distribution assumption, which is known to be invalid at least for transcriptomic data [30–32]. Importantly, the utilization of variance stabilizing transformations, such as log-scaling, would violate the linearity assumption in Equations (1)-(2) and therefore lead to biased estimation [33].

We introduce Unico, a deconvolution method for learning cell-type signals from an input of large heterogeneous bulk data and matching cell-type proportions. In practice, the latter is estimated from the input bulk profiles using reference-based decomposition (e.g., [14, 34]), as performed by all existing deconvolution methods [7–11]. The primary novelty of Unico stems from taking a model-based approach following Equation (2) while making no distributional assumptions, which renders it the first principled model-based method that is theoretically justified for analyzing cell type mixtures in any bulk genomic dataset (Methods).

A second key component of Unico is the consideration of covariance between cell types. Genomic features may be different yet coordinated across different cell types; for example, transcriptional programs can persist through multiple differentiation steps [35, 36]. Indeed, we observe that many genes present a non-trivial correlation structure across their cell-type-specific expression levels, as measured by entropy of the correlation matrix (Figure 1b), with stronger cell-type correlations (lower entropy) observed between cell types that are close in the lineage differentiation tree (Supplementary Materials). In the presence of covariance, Unico leverages the information coming from the coordination between cell types for improving deconvolution (Figure 1c,d).

### Establishing a new state-of-the-art deconvolution for bulk genomics

We compared Unico to CIBERSORTx, TCA, and bMIND, as well as to a simple baseline approach of naively weighting each bulk profile by the cell-type proportions of the sample. Our evaluation excluded methods that are either not publicly available [11] or require multiple measurements for every sample [9].

In order to form a basis for evaluation, we generated pseudo-bulk mixtures using single-cell RNAseq (scRNAseq) data from peripheral blood mononuclear cells (PBMC; n=118 donors) [29] and from lung parenchyma tissues (n=90 donors) [37] (Methods). We first evaluated the performance of the different methods in estimating population-level cell-type means, variances, and covariances by establishing gold standard estimates using the known underlying cell-type profiles of the mixtures. Our results yielded Unico, TCA, and bMIND as the best-performing methods for estimating population-level means and variances (Figure 2a; Supplementary Figures S1). Unico stands out as the leading method for learning cell-type level covariances, showcasing an average correlation improvement of 36.3% over bMIND, the second-best performing method, which also explicitly models cell-type covariance [10] (Figure 2a; Supplementary Figures S1). The ranking of methods remained consistent across different numbers of modeled cell types and various sample sizes (Supplementary Figures S2-S7).

**Figure 2:**
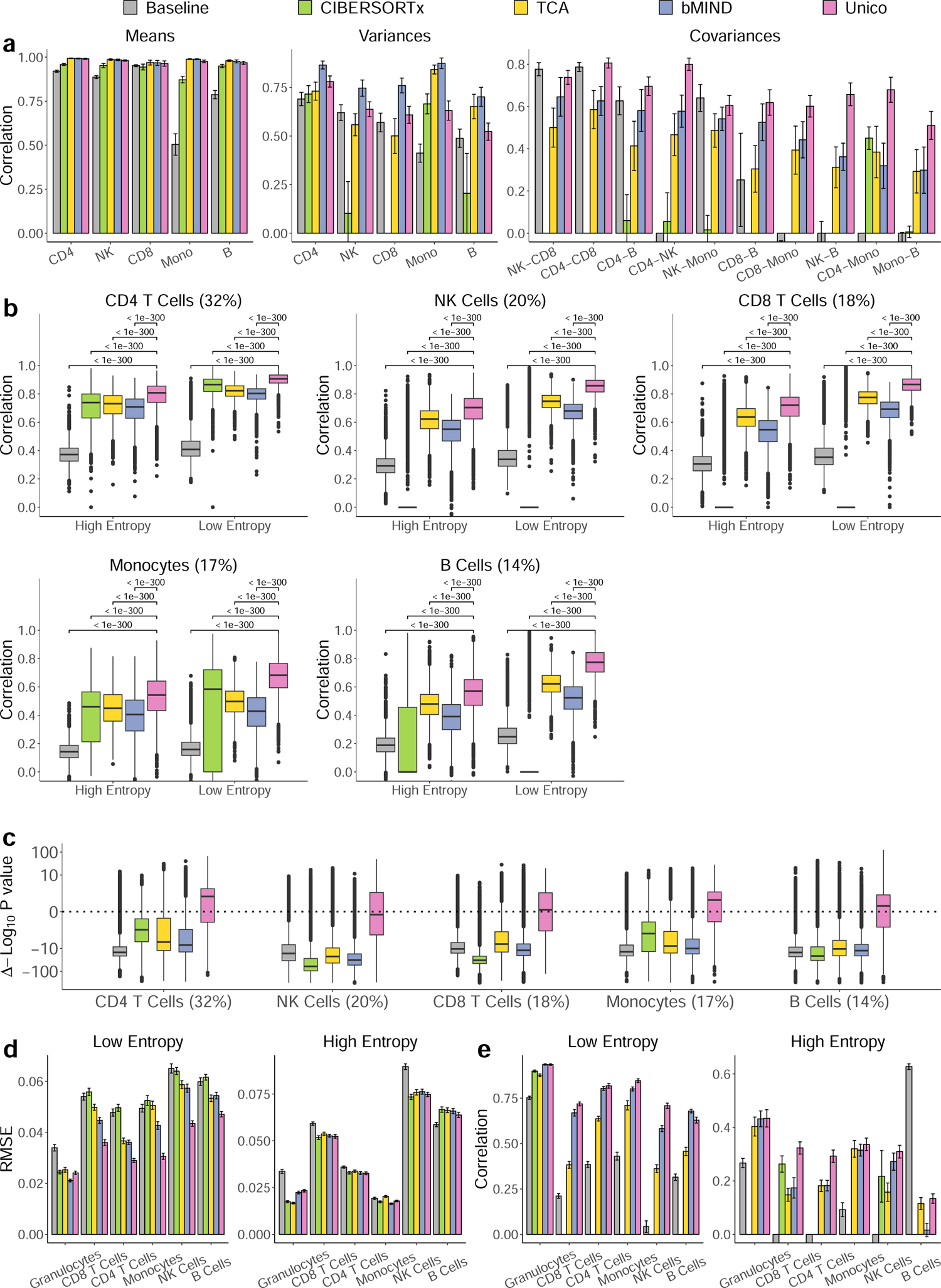
Evaluation of deconvolution methods. (a) Correlation between deconvolution and single-cell-based estimates of population-level means, variances, and covariances at the cell-type level across 20 sets of pseudo-bulk mixtures from PBMC scRNAseq profiles of five cell types (500 samples and 600 randomly selected genes in each set). (b) Evaluation of the concordance between the deconvolution estimates and the known cell-type profiles of the same data in (a). Boxplots reflect the distribution of linear correlation across all genes, and percentages indicate average cell-type abundances. (c) Assessing deconvolution estimates for their information that cannot be explained by pseudo bulk expression. Boxplots reflect the distribution across genes from the same data in (b) of Δ log_10_(p-value), the difference between the log-scaled p-values of the effects of the pseudo bulk expression and deconvolution estimates (higher is better; Methods). (d)-(e) Evaluation of whole-blood DNA methylation deconvolution in terms of RMSE and correlation between estimates and experimentally validated cell-type level methylation across 20 random sets of 1,000 highly variable CpGs. All barplots and error bars in the figure represent means and one standard deviation errors; negative correlations were truncated for visualization purposes, and p-values were calculated using a paired Wilcoxon test.

We next evaluated how well the 3D tensor estimated by Unico correlates with the true underlying cell-type expression levels of the pseudo-bulk profiles. Unico consistently outperformed the alternative methods across all cell types, providing an average improvement of 17.8% in correlation over TCA, the second-best performing method (Figure 2b; Supplementary Figures S1). Unlike Unico, bMIND is a Bayesian method that can perform deconvolution while incorporating prior information on the cell-type level means and covariates. We, therefore, further compared Unico to bMIND in the presence of informative priors from single-cell data. Remarkably, we found that bMIND could not improve upon Unico even in the unrealistic extreme case where the prior was learned from the true cell-type levels of all samples in the data (Supplementary Figures S8 and S9).

As anticipated, the improvement of Unico is more pronounced in genes that exhibit strong cell-type covariance structure (low-entropy genes; average correlation improvement of 20.0%) compared to high-entropy genes (average improvement of 14.9%). This discrepancy highlights the added information Unico gains by modeling the cell-type covariance structure. Importantly, learning a richer model does not come at the cost of significant computational runtime in this case; in fact, Unico is the second fastest deconvolution method (Supplementary Figure S10). The overall ranking of methods remained consistent across different numbers of modeled cell types and various sample sizes (Supplementary Figures S2-S7), as well as across varying levels of noise added to the cell-type proportions input (Supplementary Figures S11 and S12; Supplementary Methods).

Crucially, pseudo-bulk profiles are correlated with their true underlying cell-type levels. We therefore asked whether the 3D tensors estimated by Unico and other methods explain the variation of the true tensor beyond the pseudo-bulk input (Methods). Strikingly, we found that Unico is the only method that learns substantial variation of the true tensor when accounting for the pseudo-bulk profiles, including in lowly abundant cell types (Figure 2c; Supplementary Figures S1-S7 and S11-S12).

Lastly, we aimed to confirm the transferability of Unico to other data modalities by deconvolving bulk DNA methylation data. Reinius et al. [38] assayed from the same six individuals both whole-blood methylation and cell-type methylation of six whole-blood cell types. This data collection allowed us to establish a ground truth for the cell-type levels composing the whole-blood bulk samples. In order to circumvent the sample size limitation of the Reinius data (n=6), we took a two-step, reference-based approach. Initially, we employed Unico to estimate the model parameters using a separate large whole-blood methylation dataset from a similar population [39]. Subsequently, we utilized these parameter estimates in Unico’s tensor estimator, which given the model parameters, deconvolves the bulk profile of each individual sample independently of other samples in the data. A similar procedure was adapted for the competing methods (Methods).

Unico demonstrated exceptional performance compared to the alternative methods in reconstructing the experimentally known 3D tensor. Considering the top 10,000 most variable methylation CpGs in the data, Unico achieved an average improvement of 8.8% and 8.1% in root median squared error (RMSE) and correlation compared with bMIND, the second best performing method (Figure 2d,e; Supplementary Figures S13 and S14). The ranking of the methods was preserved when considering a set of 10,000 randomly selected CpGs; unsurprisingly, all methods present a noticeable decrease in performance in this case (Supplementary Figures S15-S17).

### Detecting cell-type-specific differential expression in heterogeneous tumors

Follicular lymphoma (FL) is the second most common indolent non-Hodgkin lymphoma (NHL) in the USA and Europe, accounting for nearly 20% of all NHL cases [40]. Previous work using FACS-sorted B cells from FL tumors identified 612 differentially expressed genes in the presence of CREBBP mutation [41]. Here, similarly to previous analysis [8], we asked whether deconvolving bulk FL tumors (n=24, including 14 with CREBBP mutation) [8, 41] would allow us to detect the previously reported effects in B cells from FL tumors. Indeed, B cell expression levels estimated by Unico from bulk FL tumors recapitulate the previously reported down- and up-regulation effects in FL B cells significantly better than alternative deconvolution methods (Figure 3a). More specifically, none of the methods performed significantly better than the others on the up-regulated genes, with the exception of the baseline method, which performed worse than all deconvolution methods. However, Unico performed best on the down-regulated genes, and remarkably, it was the only deconvolution method that performed significantly better than a straightforward bulk analysis (adjusted p-value*<*0.05; Paired Wilcoxon test).

**Figure 3:**
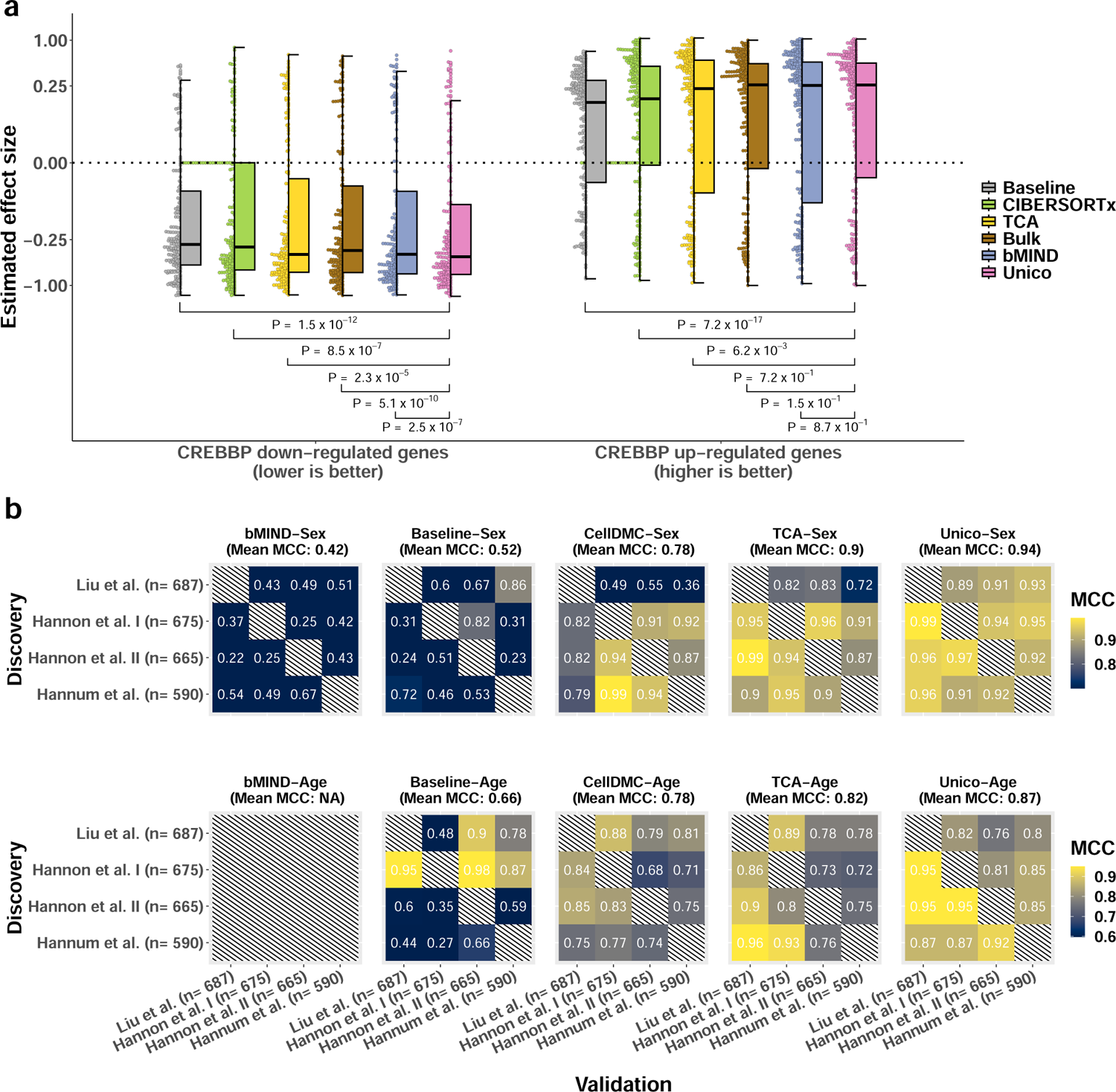
Application of deconvolution to downstream analysis tasks. (a) Deconvolution of bulk FL tumor samples for assessing previously reported CREBBP mutation-related gene expression in B cells. Presented are deconvolution-based B cell effect size distributions for 219 down-regulated and 275 up-regulated genes; comparisons to Unico were calculated using a one-sided paired Wilcoxon test. (b) Consistency in calling cell-type level differential methylation with sex and age across four independent whole-blood DNA methylation datasets. Color gradients represent the Matthews correlation coefficient (MCC) for every possible pairing of two datasets as discovery and validation (Methods). Since bMIND was designed for binary conditions only, it was not evaluated in the age analysis

### Unico improves resolution and robustness in epigenetic association studies

We expected that modeling and effectively estimating cell-type covariance would allow Unico to yield better performance in downstream applications that aim at disentangling signals between cell types. In order to demonstrate this, we evaluated the different deconvolution methods in calling cell-type level differential methylation (DM). While ground truth DM is generally unknown, one can consider the consistency of a given method across different datasets as a surrogate for true/false positive/negative rates.

We applied each method for testing a set of 177,207 CpGs for cell-type level DM in four large whole-blood methylation datasets (n*>*590 each) with sex and age information [39, 42, 43]. Specifically, for every possible combination of two out of the four datasets as discovery and validation data, we measured the consistency between datasets using the Matthews correlation coefficient (MCC) [44] (Methods). We excluded from this analysis CIBERSORTx, due to its runtime (Supplementary Figure S10) and poor performance in deconvolving bulk methylation (Figure 2e; Supplementary Figures S13-S17). Instead, we considered CellDMC, a method that was designed specifically for detecting cell-type level DM by evaluating linear effects of interaction terms between the condition of interest and cell-type proportions [45]. We observe that Unico provides the best overall consistency (Figure 3b), and it significantly improves upon TCA, the second best method (p-value*≤*0.05 for both sex and age; one-sided paired Wilcoxon test). Importantly, the runtime of Unico was on par with TCA’s (Supplementary Figure S10).

The above evaluation disregards a straightforward analysis of the bulk data, which cannot associate DM with specific cell types but rather call CpGs as generally associated with conditions (“tissue-level” analysis). Intuitively, models that provide cell-type resolution are more realistic and are thus expected to improve cross-dataset consistency over a standard tissue-level analysis. In order to verify this intuition, we evaluated a standard linear regression analysis of the bulk data for calling tissue-level DM (Supplementary Figure S18). We observe that cell-type level analysis using any of the deconvolution methods provides a substantial improvement in consistency compared to the bulk analysis. In particular, Unico provides an increase of 107.5% and 40.7% in MCC for sex and age, respectively. Further adapting the different deconvolution methods to call tissue-level DM (Supplementary Methods) yields all methods as better than a standard bulk analysis, with Unico being the top performing method (Supplementary Figure S18) These results demonstrate how carefully modeling the cell-type signals in bulk data improves analysis even if constrained to a tissue-level context.

## 3 Discussion

We propose Unico, a deconvolution method that is theoretically appropriate for any bulk genomic data type that reflects mixtures of signals across cell types. Here, we demonstrate the utility of Unico for gene expression and DNA methylation, however, our distribution-free treatment suggests its applicability to other genomic data types as well. Unico leverages covariance across cell types, and as such, it deconvolves particularly well low-entropy features that exhibit non-trivial correlation structures between cell types. Remarkably, our evaluation, based on two scRNAseq datasets from different tissues and purified methylation data, demonstrates that Unico considerably outperforms state-of-the-art methods in general, even when deconvolving high entropy features.

Our deconvolution of pseudo-bulk RNA mixtures benchmarked the performance of several deconvolution methods. Notably, we did not evaluate the correlation between the cell-type expression and the bulk levels in this analysis. Instead, we showed that Unico is the deconvolution method that provides the most additional cell-type information beyond the bulk levels. At least in some cases, bulk expression is expected to be more correlated with cell-type expression; in particular, bulk levels are expected to yield an almost perfect correlation for genes presenting the same expression patterns across cell types. Yet, we omit an evaluation of standard bulk levels in this specific analysis and consider them only in downstream analysis. A key reason for that stems from the need to compare bulk and deconvolution estimates on equal ground. This requires dissecting and mapping the variation in bulk levels to different cell types to match in resolution to a cell-type deconvolution output. An arguably natural approach to achieve this is weighting the bulk levels by cell-type proportions, which we included as a baseline model throughout our study.

Another key reason for considering standard bulk levels only in the context of downstream analysis relates to the core motivation of performing deconvolution. The premise of deconvolution is to provide novel biological insights by modeling and differentiating cell-type patterns in the context of conditions. Thus, the utility of deconvolution should ultimately be established by advancing upon bulk-based downstream analysis tasks. Under this view, evaluating the correlation of deconvolution estimates with cell-type patterns merely represents an indirect (yet common) metric for comparing deconvolution methods. Such correlations rely on point estimates of the cell-type patterns, which are expected to be noisy, and reflect the “best guess” of a given model. Yet, deconvolution methods such as Unico also model the uncertainty of those point estimates. This uncertainty can be integrated to perform well-powered downstream statistical analysis without explicitly generating noisy point estimates. As we demonstrate through the analysis of multiple RNA and DNA methylation datasets, this approach advances over a standard bulk analysis in terms of standard performance metrics and its ability to report results at a cell-type resolution.

Finally, Unico has some limitations, and while these limitations are not unique to Unico but are rather common to all the deconvolution methods we evaluated, they may potentially bias and affect the performance of our proposed model. First, Unico makes the assumption that cell-type proportions of the input bulk samples are known. Admittedly, this information is rarely available in bulk genomics data, so proportions need to be estimated in practice. While it is commonplace to employ reference-based methods for learning cell-type compositions, using estimates in place of measurements creates a source of noise and potential bias. Our different real data analyses using precisely such estimates, as well as our evaluation of deconvolution of in-silico mixtures under noisy cell-type compositions, suggest that, in practice, Unico is overall robust to this source of noise.

Other limitations of Unico pertain to the fact that lowly abundant cell types represent only a small fraction of the variance in bulk data. Unico is, therefore, expected to perform poorly when attempting to model a large number of cell types. Since heterogeneous tissues often represent mixtures of a large number of cell types and subtypes, the deconvolution of Unico may be biased by unmodeled cell types. Despite these concerns, we conclude that our comprehensive evaluation of Unico across diverse datasets and data modalities provides compelling evidence of its superiority over existing state-of-the-art deconvolution methods.

## 4 Methods

### Unico: a model for uniform cross-omics deconvolution

We denote *X_ij_* to be the (tissue-level) bulk gene expression in sample *i ∈ {*1*…, n}* of gene *j ∈ {*1*…, m}*. For simplicity of exposition, we use the notion of gene expression, however, *j* can represent any other genomic feature that may vary across cell types. We assume:

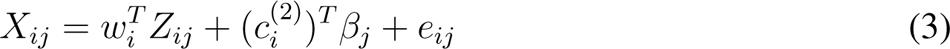

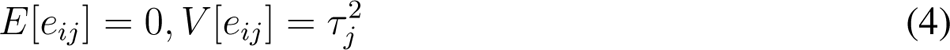

The first term in Equation (3) defines *X_ij_* as a weighted linear combination of cell-type expression levels. Specifically, *w_i_* = (*w_i_*_1_*, …, w_ik_*) is a vector of sample-specific cell-type proportions of *k* cell types that are assumed to compose the studied tissue and *Z_ij_* = (*Z_ij_*_1_*, …, Z_ijk_*) is a vector of cell-type expression levels of gene *j* in sample *i*. The second and third terms in Equation (3) model systematic and non-systematic variation. Specifically, *e_ij_* is an i.i.d. component of variation that reflects measurement noise, *c*^(2)^ is a *p*_2_-length vector of known covariate values of sample *i* that may be associated with unwanted global effects (i.e., “tissue-level” effects that may affect many genes and are not cell-type-specific, such as batch effects), and *β_j_* is a vector of the corresponding gene-specific fixed effect sizes.

We assume that cell-type proportions *{w_i_}* are fixed and given. In practice, these can be estimated using a reference-based approach (e.g., [14, 34]), as suggested by other deconvolution methods [7–11]). In contrast to a standard decomposition problem, which assumes shared cell-type expression levels across all samples, the unknown *{Z_ij_}* components are modeled as random variables; this is emphasized by the use of upper-case notation. Specifically, for *Z_ijh_*, the gene expression in sample *i* of gene *j* and cell type *h ∈ {*1*…, k}*, we assume:

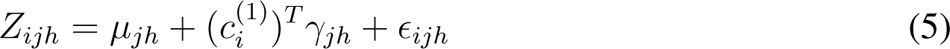

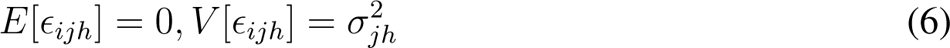

where *µ_jh_* is the mean level, specific to gene *j* and cell type *h*, *ɛ_ihj_* is an i.i.d. noise term with mean zero and variance *σ*^2^ that may be specific to gene *j* and cell type *h*, *c*^(1)^ is a *p*_1_-length vector of known covariate values of sample *i* that may present cell-type-specific effects, and *γ_jh_*is a vector of corresponding fixed effect sizes.

Lastly, we further model cell-type covariance. Concretely, we model the covariance of a given gene *j* across cell types *h, q* and denote:

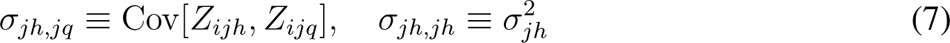

The Unico model makes no assumptions on the distribution of the components of variation in Equations (3) and (5), which makes it naturally applicable to all heterogeneous tissue-level omics that can be represented as linear combinations of cell-type level signals. Finally, Unico can be viewed as a generalization of the TCA model and as a frequentist alternative for the bMIND model. See Supplementary Methods for details.

### Estimating the underlying 3D tensor with Unico

Given a single realization *x_ij_* of the bulk level coming from *X_ij_*, we wish to learn *z_ij_*, the realization of the cell-type-specific expression levels *Z_ij_* of the corresponding sample *i* and gene *j*. Our goal is hence to compose a 3D tensor (samples by genes by cell types) based on the 2D input matrix. We address this problem by setting the estimator of *z_ij_* to be the expected value of the conditional distribution *Z_ij_|X_ij_*:

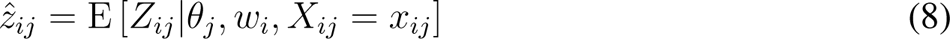

where *θ_j_* is the set of parameters that are specific to gene *j*, that is,

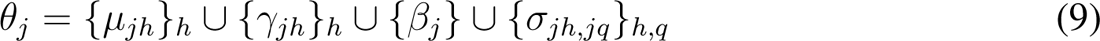

The following theorem provides an analytical solution for the estimator *z*^*_ij_*under the Unico model in Equations (3)-(7).

### Theorem 1 (The Unico 3D tensor estimator)

The solution for the estimator stated in Equation (8) under the Unico model is given by:

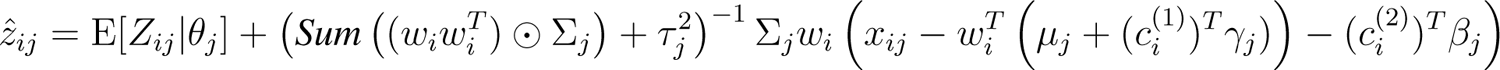

where 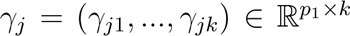 is a martix composed of the vectors 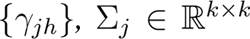 is the cell-type covariance matrix of gene j, the ⊙ operator is the Hadamard product of two matrices, and the Sum(·) operator is a summation across all entries of a matrix.

Proof is given in the Supplementary Methods.

Theorem 1 provides an analytical solution for the 3D tensor given the cell-type proportions *{w_i_}* and model parameters *θ_j_*. As mentioned above, in practice, cell-type proportions are estimated using decomposition methods, and as we later describe, the model parameters can be estimated from the observed bulk data and the estimated cell-type proportions.

Unico essentially defines the estimator *z*^*_ij_* as the expected value of the conditional distribution *Z_ij_|X_ij_* = *x_ij_*, which was previously suggested in TCA [7]. However, under the richer Unico model, this conditional distribution becomes more informative owing to the correlation structure between cell types. Intuitively, learning cell-type levels that better capture cell-type covariance will enhance our capacity to assign deconvolution signals accurately to the respective cell types in downstream analysis.

A-priori one may wonder whether modeling cell-type covariance is necessary for a deconvolution method to recapitulate the true cell-type covariance in the data. Put differently, one could expect an accurate deconvolution method to capture cell-type covariance regardless of an explicit modeling of the covariance. However, our empirical results suggest that such modeling is valuable for accurate deconvolution, and the following theorem provides intuition into why modeling the covariance is indeed desired in order to achieve accurate deconvolution. Besides Unico, TCA [7] is the only existing deconvolution method that offers an analytical estimator for the 3D tensor. Hence, the following exclusively focuses on Unico and TCA, as the theoretical analysis for other methods remains unclear.

### Theorem 2 (Improved capacity to reduce covariance bias)

*Assume for simplicity ∀h*: *µ_jh_* = 0, *σ*^2^ = 1*, τ_j_* = 0*, and no covariates for some feature j under Equations (3)-(7). If n → ∞ then* (i) *the cell-type covariances of the 3D tensor estimated by TCA are fixed and do not depend on feature j, and (ii) the cell-type covariances of the 3D tensor estimated by Unico are a function of the cell-type covariance of feature j*. Proof is given in the Supplementary Methods.

### Optimization

We estimate the parameters of the model by following concepts from the Generalized Method of Moments (GMM) [46]. The GMM framework allows us to learn the parameters of a model by iteratively solving equations (moment conditions) that match population moments (or, more generally, a function of population moments) with their corresponding data-derived sample moments. We tailor the optimization to the Unico model to form asymptotically consistent estimators as in classical GMMs [46], while introducing practical considerations and constraints that are essential for finite data. The full details about the optimization and implementation of Unico are provided in the Supplementary Methods.

### Implementation of Unico and practical considerations

We implemented Unico in R. In order to stabilize the parameter estimation, in practice, we consider non-negativity constraints when estimating the means and a small *L*_2_ penalty when estimating the variances and covariances in the model. The latter alleviates the risk of multicollinearity and, therefore, inaccurate estimation owing to the high correlation between the proportions of different cell types. Additionally, when estimating the parameters of a given feature, we disregard samples with values that diverge from the mean by more than two standard deviations. This measure prevents extreme and non-representative data points from dominating the solution.

We optimize the Unico model iteratively. At the end of each iteration, we update the weights, which can then be used for weighting the samples in the following iteration (Supplementary Methods). At a given iteration, we learn the means using the constrained least squares solver `pcls` from the `mgcv` R package, and we learn the variances and covariances using the COBYLA algorithm [47] as implemented in the `nloptr` R package [48]. Empirically, we found that Unico works well using as few as two iterations (i.e., updating the weights once) for estimating the means and using three iterations for estimating the variances and covariances (data not shown).

### PBMC and lung scRNAseq data

We obtained the PBMC scRNAseq dataset from a COVID-19 study by Stephenson et al. [29]. We arbitrarily selected only one sample for donors with multiple measurements, which resulted in a total of 118 samples for the analysis. After excluding cells with a high percentage of hemoglobin (*≥* 1%) or mitochondria (*≥* 5%), and low percentage of ribosomal content (*≤* 1%), in addition to requiring a minimal and maximal number of unique expressed genes (*≥* 500*, ≤* 2500) and total UMI counts (*≥* 2000*, ≤* 15000), 499,336 cells remained for the analysis. In addition, we used scRNAseq from the data collection presented by Sikkema et al. [37] as part of a study for integrating multiple datasets collected from the human respiratory system. We focused on the lung parenchyma samples (n=90) that composed most of the carefully annotated group of samples in the original study (defined by the authors as the “core reference” group). Employing the same data filtering criteria as for the PBMC data resulted in a total of 296,227 cells for the analysis. For both the PBMC and lung datasets we used the cell-type annotations provided by the authors and applied a counts per million (CPM) normalization.

### Gene expression data with follicular lymphoma

We used a preprocessed microarray bulk FL data (n=302) by Newman et al. [8]. In total, out of the 302 samples available, 14 were confirmed to have the CREBBP mutation, and 10 samples were confirmed to exhibit a wild-type allele. The CREBBP status for 12 of these samples was collected by Green et al. [41] and the remaining 12 samples by Newman et al. [8]; the CREBBP status of all 24 samples was made available in the supplementary files of Newman et al. For defining a ground truth list of differentially expressed genes with CREBBP mutation in FL B cells, we considered the set of 334 up-regulated and 279 down-regulated genes that were previously reported in a study with sorted B cells from FL tumors [41]. Intersecting these sets with the genes available in the bulk FL data left us with 275 and 219 up- and down-regulated genes for evaluation.

### Whole-blood DNA methylation datasets

We used a total of five beta-normalized DNA methylation datasets that were collected using the Illumina 450K methylation array. For the methylation deconvolution analysis, we obtained data from Reinius et al. [38], including whole-blood (n=6) and matching cell-sorted methylation data from the same individuals (granulocytes, monocytes, NK, B, CD4 T, and CD8 T cells). For the cell-type level differential methylation (DM) analysis, we considered whole-blood datasets from Liu et al. (n=687) [42], Hannum et al. (n=590; samples with missing smoking status were excluded) [39], and two datasets from Hannon et al. (n=675, n=665) [43]. In all datasets, we removed CpGs with non-autosomal, polymorphic, and cross-reactive probes [49], and we excluded low variance CpGs (variance*<*0.001). This left us with 153,155, 144,743, 134,250, and 95,360 CpGs for the Liu, Hannum, and the two Hannon datasets, respectively. For the Reinius dataset, we considered CpGs at the intersection between the Reinius data and a preprocessed version of the Hannum dataset (restricted to samples with European ancestry; 93,086 CpGs). Lastly, cell-type proportions were estimated for all whole-blood datasets using EpiDISH, a reference-based methylation decomposition method [50].

### Implementation and application of competing deconvolution and cell-type association methods

We ran all CIBERSORTx [8] related codes under a docker container version 1.0 encapsulating both the “High Resolution” mode (for estimating cell-type level profiles) and the “Fractions” mode (for estimating cell-type proportions) with default parameters and authentication token granted by the CIBERSORTx team upon request. CIBERSORTx evaluates the maximum value in a bulk input and automatically assumes the data have been log-normalized if the maximum is less than 50. This choice is reasonable for transcriptomic data, for which CIBERSORTx was designed, however, it is not justified for beta-normalized methylation levels that are restricted to the interval [0, 1]. We thus scaled the methylation beta values by a factor of 10,000 prior to the application of CIBERSORTx and rescaled the results back to original scale.

We installed the TCA [7] R CRAN package version v1.2.1 deposited on CRAN and evaluated its performance under default parameters. We fitted the model using the function `tca` and performed deconvolution using the `tensor` function. For the cell-type level DM analysis, both the joint (tissue-level) and marginal (cell-type level) statistical tests were automatically evaluated as part of the model parameter learning step in the `tca` function.

bMIND [10] is available via the MIND R CRAN package version 0.3.3. We obtained the cell-type specific profiles and the estimated model parameters with the function `bMIND` and performed association testing with the function `test`. bMIND evaluates the maximum value in the bulk input and automatically log transforms the data if the maximum is larger than 50. We, therefore, scaled the bulk expression profile (and the single-cell derived prior) by the inverse of the largest detected value before applying bMIND, and then rescaled the output back to the original scale. This approach ensured consistency and comparability across all deconvolution methods. Specifically, allowing the default log transformation of the data would have violated the assumption that bulk levels represent linear combinations of cell-type levels.

Throughout this work, we also evaluated a baseline approach in our analysis and evaluation by simply considering the product of the observed bulk data and the cell-type proportions as cell-type level estimates. That is, we estimated *z_ijh_*, the cell-type level of sample *i*, gene *j*, and cell type *h* as *z*^Baseline^ = *x_ij_ · w_ih_*. Finally, we applied CellDMC [45] for DM using the implementation in the Bioconductor R package EpiDISH, version 2.10.0.

### Deconvolving mixtures of gene expression profiles and estimating cell-type level moments

We used both the PBMC and lung scRNAseq datasets for generating pseudo-bulk mixtures. Briefly, for creating a new pseudo-bulk sample, we first drew (with replacement) all cell-type level profiles of one randomly selected sample. The cell-type profiles of each individual sample were defined as normalized pseudo-bulk counts at the cell-type level. We then drew (with replacement) the cell-type proportions of one randomly selected sample in the data (total number of cells coming from each cell type, normalized to sum up to 1). Eventually, these were used as the weights for a weighted linear combination of the cell-type level profiles to create one pseudo-bulk sample.

In the PBMC analysis, we considered either five major cell-type groups (monocytes, NK, B, CD4 T, and CD8 T cells) or seven cell types by further stratifying B cells into canonical B cells and plasma cells and monocytes into CD16 and CD14 monocytes. In the analysis with lung cells, we considered either four major cell-type groups (endothelial, stromal, immune, and epithelial cells) or six cell types by further stratifying immune cells into myeloid and lymphoid compartments and epithelial cells into the airway and alveolar epithelium cells. Our evaluation was restricted to the top 10,000 most highly expressed genes in the data. See Supplementary Methods for more details.

The pseudo-bulk mixtures, along with the corresponding mixing proportions, were provided as the input for all deconvolution methods to learn 3D tensors. We assessed these tensors for their accuracy by comparing them against the known cell-type profiles. Particularly, for a given cell type and a given gene, we evaluated the correlation between the true cell-type expression levels of the pseudo-bulk samples and their deconvolution-based estimates.

We obtained estimates of population-level cell-type moments from the data (means, variances, and covariances per gene) directly from the output of the deconvolution methods. For methods that do not explicitly output such estimates (e.g., no method except for bMIND and Unico outputs covariance estimates), we used the estimated tensor for calculating these moments. To evaluate the accuracy of the estimated moments, we established gold standard estimates based on the cell-type profiles underlying the pseudo-bulk mixtures. In order to mitigate the potential influence of outliers, we considered only samples within 2 standard deviations from the mean for the estimation of moments of a given gene.

Finally, we used multiple linear regression to evaluate whether an estimated 3D tensor of a given deconvolution method captures the variation of the true tensor beyond its correlation with the deconvolution input (i.e., pseudo-bulk and cell-type proportions). In more detail, for every gene and cell type, we fitted a regression model for the known cell-type expression levels as the dependent variable using several independent variables, including the pseudo-bulk levels of the gene, the cell-type proportions, and the cell-type tensor estimates. This allowed us to quantify to what extent the deconvolution-based estimates provide information beyond the bulk data. Specifically, we defined Δ log_10_(p-value) as the difference between the log-scaled (basis 10) t-test derived p-values of the pseudo-bulk variable and the estimated cell-type levels in the regression. Of note, we defined the p-values to be 1 in cases where cell-type levels were estimated to have no variation. In order to mitigate potential biases due to heavy-tailed distributions of expression levels, we log1p-transformed expression levels and considered only samples within 2 standard deviations from the mean.

### Deconvolving the Reinius whole-blood DNA methylation data

Unlike our deconvolution of gene expression mixtures, the size of the Reinis data (n=6) does not allow for drawing reliable conclusions through a straightforward evaluation. Particularly, Unico, as well as current deconvolution methods, are designed to operate on large bulk data. We circumvented this limitation by taking a two-step reference-based procedure. First, we learned the parameters of the Unico model from the larger Hannum whole-blood methylation data [39]. Acknowledging that population structure affects methylation [51], we focused solely on Caucasian individuals from the Hannum data (n=426), anticipating that they would adequately represent the Swedish individuals in the Reinius study. Then, we plugged these parameter estimates into Unico’s 3D tensor estimator together with the Reinius bulk profiles and their cell-type proportion estimates. We performed the same procedure for TCA, however, CIBERSORTx and bMIND, which do not provide an analytical estimator of the tensor, required a different strategy. In order to inform the deconvolution of CIBERSORTx and bMIND with the same additional information, we applied these methods to the concatenation of the Reinius and Hannum datasets and extracted the cell-type level estimates for the Reinius samples.

Benchmarking methods based on the Reinius data presents a second challenge: determining a proper way to evaluate their performance given that data from only six individuals is available for the analysis. We tackle this limitation by collapsing methylation levels in the estimated tensor along both the CpG and sample axes. That is, for every cell type, we evaluated how correlated is the vector of all methylation estimates of the cell type (i.e., by pooling estimates across all CpGs and samples) with the experimentally measured ground truth levels from purified cells. This yielded a single correlation score per cell type. Importantly, when stacking CpGs for evaluation, a deconvolution that only correctly estimates relative means and scales of CpGs but performs poorly in terms of per-CpG correlation (i.e., across samples) may achieve spuriously high correlation levels. We addressed this by removing from every CpG its cell-type level mean methylation level.

Since beta-normalized methylation levels are bounded to the range [0,1], unlike in the deconvolution of relative expression levels, we further evaluated the divergence of the estimated 3D tensors from the true cell-type levels in absolute terms. Specifically, we evaluated the root median square error (RMSE) between the true and each estimated 3D tensor; we expected that a median metric in place of a standard mean square error would improve robustness to outliers. Similarly to the evaluation of correlation, we calculated an RMSE value per cell type after collapsing methylation levels in the tensors along both the CpGs and samples axes.

Finally, our benchmarking focused either on randomly selected CpGs or on a set of highly variable CpGs based on the Reinius data. For defining the latter, we ranked the CpGs in the intersection of the Reinius and Hannum datasets (93,086 CpGs) by the sum of their variances in the different cell types using the sorted methylation Reinius data and chose the top 10,000 CpGs with the largest values.

### Calculating robust linear correlation

All the correlation values reported throughout our analysis and evaluation were calculated using a robust linear correlation metric in place of the standard Pearson correlation. Specifically, we used the function `cov.rob` from the MASS R package [52], which performs an approximate search for a subset of the observations to exclude such that a Gaussian confidence ellipsoid is minimized in volume. Effectively, this procedure trims outliers that may otherwise dramatically bias correlation levels (Supplementary Figure S19). In particular, if either input vector has an interquartile range (IQR) of 0, `cov.rob` defines the correlation as 0. Throughout the paper, we set the fraction of outliers to exclude to 5% of the data points.

### Calculating von Neumann entropy

We quantify the amount of signal coming from the covariance structure of a given gene by the von Neumann entropy [53]. For a given gene, the von Neumann entropy is defined as the entropy applied to the eigenvalues of the normalized cell-type covariance matrix of the gene (i.e., a *k×k* matrix of correlations between cell types). High entropy corresponds to cases where no substantial cell-type covariance structure exists, and low entropy indicates strong positive or negative correlations between cell types. Throughout our evaluation of the deconvolution results, we grouped genes into high- and low-entropy groups. This classification was based on ranking the genes by their entropy and assigning genes with above-median (below-median) entropy to the high (low) entropy group. Lastly, the normalized von Neumann entropy presented in figure 1b simply refers to von Neumann entropy values scaled to the range [0,1]. Since the von Neumann entropy is bounded by a number that depends on the number of cell types *k*, this normalization enables us to evaluate and visualize the distribution of entropy across genes using covariance matrices of different sizes.

### Deconvolving bulk profiles from follicular lymphoma tumors

For every deconvolution method, we first estimated the 3D tensor of the bulk FL dataset (n=302) while considering only the sets of 275 and 219 genes that were previously reported as up- and down-regulated with the CREBBP mutation. We provided each method with cell-type proportions estimated using CIBERSORTx (“Fractions” mode) with the LM22 signature matrix [54], while collapsing the estimated proportions into 4 categories: B cells, CD4 T cells, CD8 T cells, and “remaining”.

A straightforward evaluation would include calculating for every method log-fold changes (LFCs) with the CREBBP mutation based on the estimated B cell expression levels. This would allow for assessing the concordance between the LFCs and the previously reported direction of the differentially expressed genes. However, the group of CREBBP-mutated tumors presents an elevated B cell composition, which is expected to lead to an overly optimistic performance on the set of up-regulated genes in cases of deconvolution estimates that are biased by cell composition (Supplementary Figure S20). Most notably, since the baseline method estimates B cell expression levels by naively multiplying bulk levels by B cell proportions, the baseline estimates are expected to be artificially higher for samples with higher B cell composition. The baseline method, therefore, consistently estimates higher B cell expression levels for the CREBBP-mutated tumors, regardless of whether the genes are truly down- or up-regulated. Consequently, genes that are truly up-regulated in CREBBP tumors are expected to present strong LFCs under the baseline given the combination of both real and artificial up-regulation effects.

In order to account for the B cell composition bias, we used multiple linear regression to test whether the estimated tensors capture the mutation effects beyond the effect of B cell composition. In more detail, for every gene, we fitted a regression model for the estimated B cell expression levels as the dependent variable using the B cell composition and the mutation status as independent variables. We performed the same procedure while using the bulk expression levels as the dependent variable to evaluate a standard analysis of bulk expression. In order to allow a comparable evaluation of the estimated mutation effect sizes across the different methods and to alleviate the potential effect of outliers, we standardized the log1p-scaled B cell expression estimates of every gene. For methods that do not constrain non-negativity in their estimated tensor, for every gene and cell type, we shifted the distribution of the estimates by subtracting the minimum value detected, which enforced non-negativity prior to the log1p transformation. The effect size of a gene that was estimated to have a constant B cell expression level across all samples was set to 0.

### Cell-type level epigenetic association studies with sex and age

We performed statistical testing for calling DM using Unico, TCA [7], bMIND [10], and CellDMC [45] (Supplementary Methods). As a baseline model, we evaluated the linear effects of the conditions on the tensor estimates of our baseline deconvolution. Concretely, for a given CpG and cell type, we fitted a linear regression model with the baseline-estimated cell-type level methylation as the dependent variable and the condition (and covariates) as the independent variable. This allowed us to calculate t-statistics and derive p-values for the cell-type level effects of the conditions under a baseline deconvolution.

Our analysis included cell-type level covariates (*{c*^(1)^*}* under the Unico notations) and tissue-level covariates (*{c*^(2)^*}* under the Unico notations). For cell-type covariates, we considered age and sex in the analysis of all four whole-blood methylation datasets (Liu et al. [42], Hannum et al. [39], and two cohors by Hannon et al. [43]). In addition, we accounted for rheumatoid arthritis and smoking status in the Liu data, schizophrenia status in the Hannon data, and ethnicity and smoking status in the Hannum data. Across all datasets, smoking status was classified into three major categories: never, past, and current smoker. For tissue-level covariates, we considered surrogates of technical variability. In more detail, for each methylation dataset, prior to filtering any CpG, we took a previously suggested approach [7, 28] of estimating factors of technical variation by calculating the top 20 principal components (PCs) of the 10,000 least variable CpGs of each methylation array. We expected these PCs to capture only global technical variation and no biological variation due to the use of CpGs with nearly constant variance. In addition to these PCs, we further accounted for plate information, which was available for the Hannum data. All the benchmarked methods were designed to account for cell-type and tissue-level covariates, except for the baseline model. For the latter, we simply included the full set of covariates as independent variables in the linear regression model.

The inter-individual distribution of array-probed methylation levels is approximately normally distributed for most CpGs. For that reason, TCA and CellDMC, which were designed for methylation data, assume the data is normally distributed; bMIND assumes normality as well, even though it was not designed for methylation. We, therefore, similarly applied statistical testing under a normality assumption when evaluating Unico on calling DM (Supplementary Methods). Notably, this assumption is not required given that the Unico framework is generally distribution-free and allows us to derive asymptotic p-values (Supplementary Methods). Indeed, we empirically observe that asymptotically-derived p-values are highly correlated with their parametric counterparts, while also being calibrated under the null (Supplementary Figure S21-S24).

For any given ordered pair of datasets (discovery and validation), we considered the CpGs at the intersection of the two datasets. True positives (TPs) were defined as CpGs that are (i) genome-wide significant in the discovery dataset under a Bonferroni-corrected threshold and (ii) significant in the validation dataset, under a Bonferroni-corrected threshold adjusting for the number of significant hits identified in the discovery data. CpGs that only satisfied condition (i) and either failed to satisfy condition (ii) or demonstrated inconsistent direction of their estimated effect size were considered false negatives (FNs). CpGs with p-value*>*0.95 in the discovery dataset were considered as negative controls for the evaluation of false positives (FPs) and true negatives (TNs). That is, negative controls with significant (non-significant) p-values under a Bonferroni-corrected threshold adjusting for the number of negative controls in the validation data were counted as FPs (TNs).

Finally, as a metric of consistency across datasets, we calculated the MCC per method for every pair of discovery and validation datasets. We favored MCC over the widely-used F1 score since the former incorporates true negatives, which makes it a better choice for assessing model performance on imbalanced class distributions [44]. Yet, for completeness, we further considered the F1 score as the consistency metric, which revealed qualitatively similar results (Supplementary Figure S25 and S26).

## Supporting information

Unico Supplementary

## Data availability

The bulk FL data is available from GEO (accession number GSE127462). The whole-blood methylation data with matching sorted cells, as well as the whole-blood methylation datasets used for cell-type level DM analysis are available from GEO (accessions GSE35069, GSE42861 GSE40279, GSE80417, GSE84727). The PBMC scRNAseq dataset was downloaded from EMBL-EBI (accession E-MTAB-10026), and the lung scRNAseq is available on cellxgene [55] as the integrated Human Lung Cell Atlas.

## Code availability

Unico is available as an R package on CRAN and GitHub under the GPL-3 license at: https://github.com/cozygene/Unico.

