## Supplementary material for "A unified model for cell-type resolution genomics from heterogeneous omics data": Unico Supplementary

### S1 Supplementary Figures

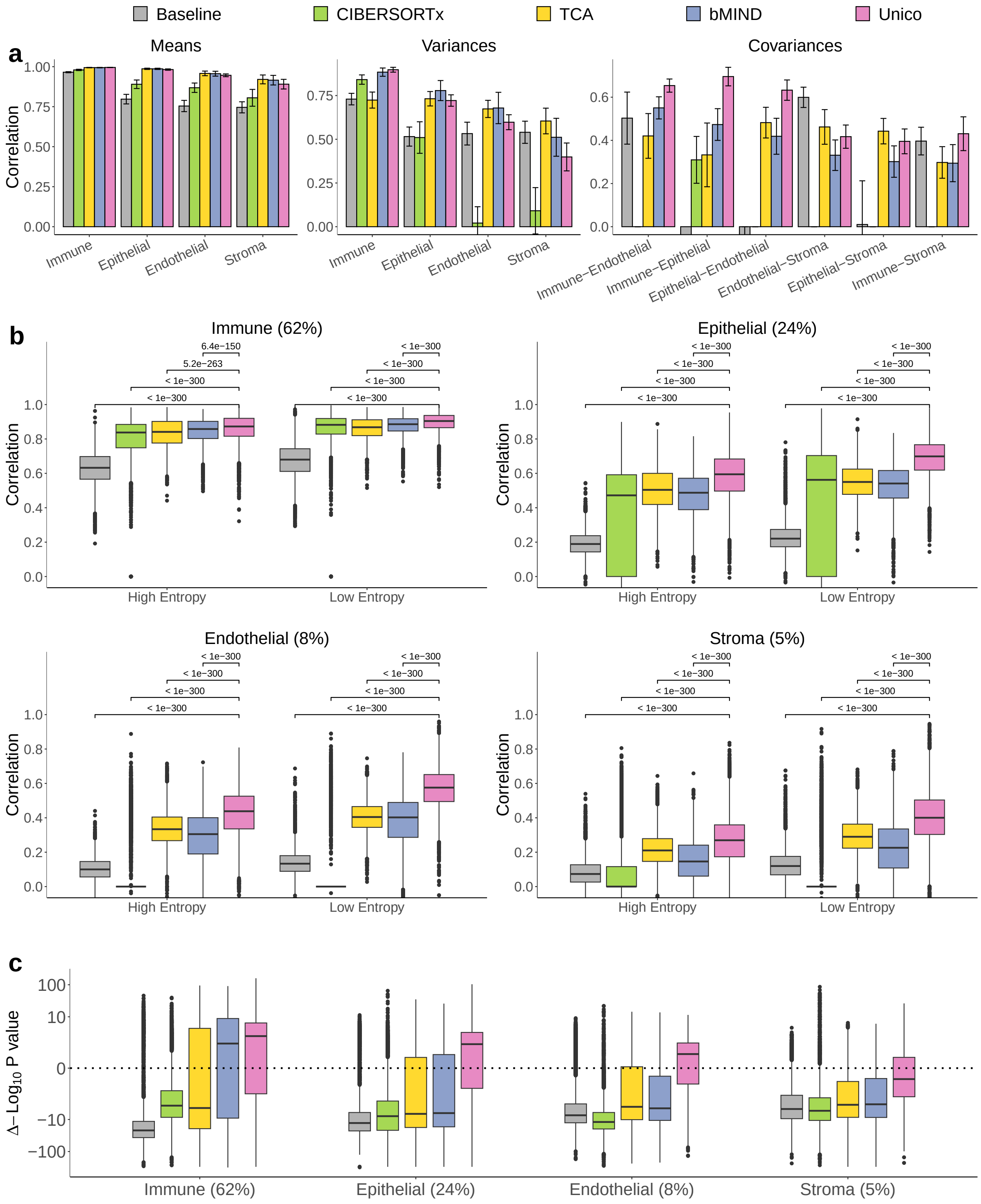

Figure S1: Evaluation of deconvolution methods on RNA pseudo-bulk mixtures. (a-c) Same analyses as in main Figure 2a-c, only using pseudo-bulk mixtures from lung scRNAseq profiles of four cell types (500 samples and 600 genes in each set)

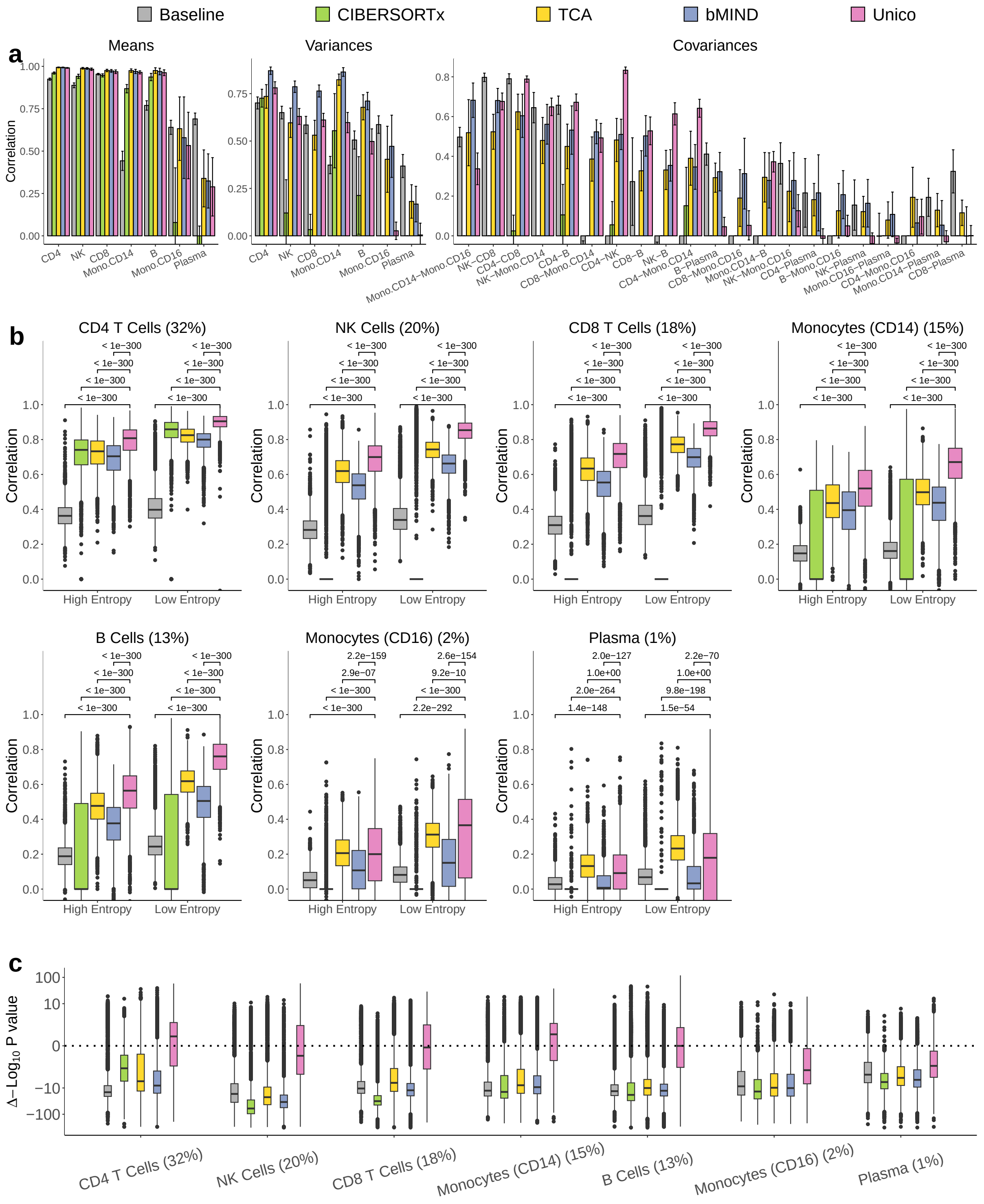

Figure S2: Evaluation of deconvolution methods on RNA pseudo-bulk mixtures. (a-c) Same analyses as in Supplementary Figure S1, but using pseudo-bulk mixtures from PBMC scRNAseq profiles of seven cell types (500 samples and 600 genes in each set)

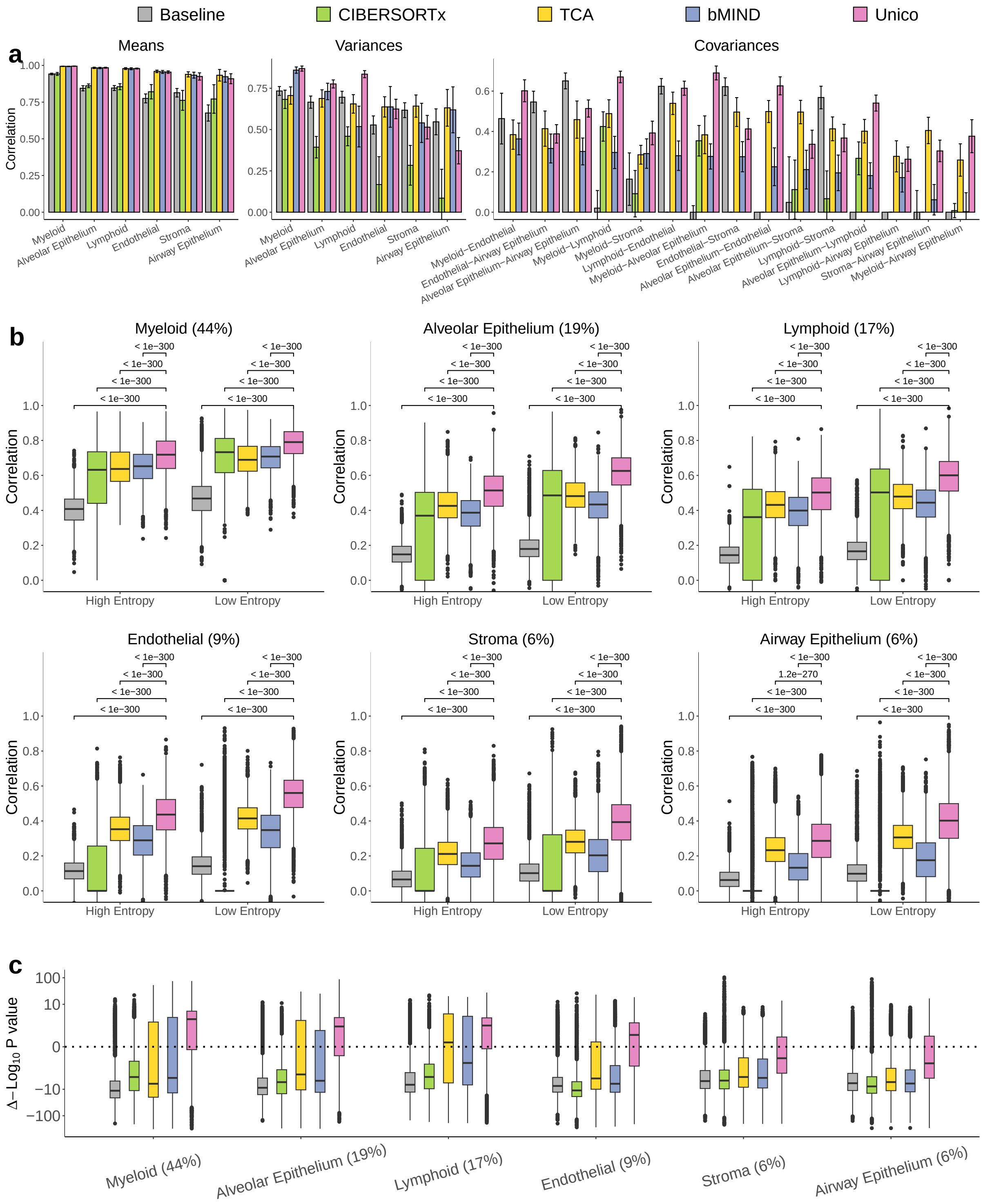

Figure S3: Evaluation of deconvolution methods on RNA pseudo-bulk mixtures. (a-c) Same analyses as in Supplementary Figure S1, but using pseudo-bulk mixtures from lung scRNAseq profiles of six cell types (500 samples and 600 genes in each set)

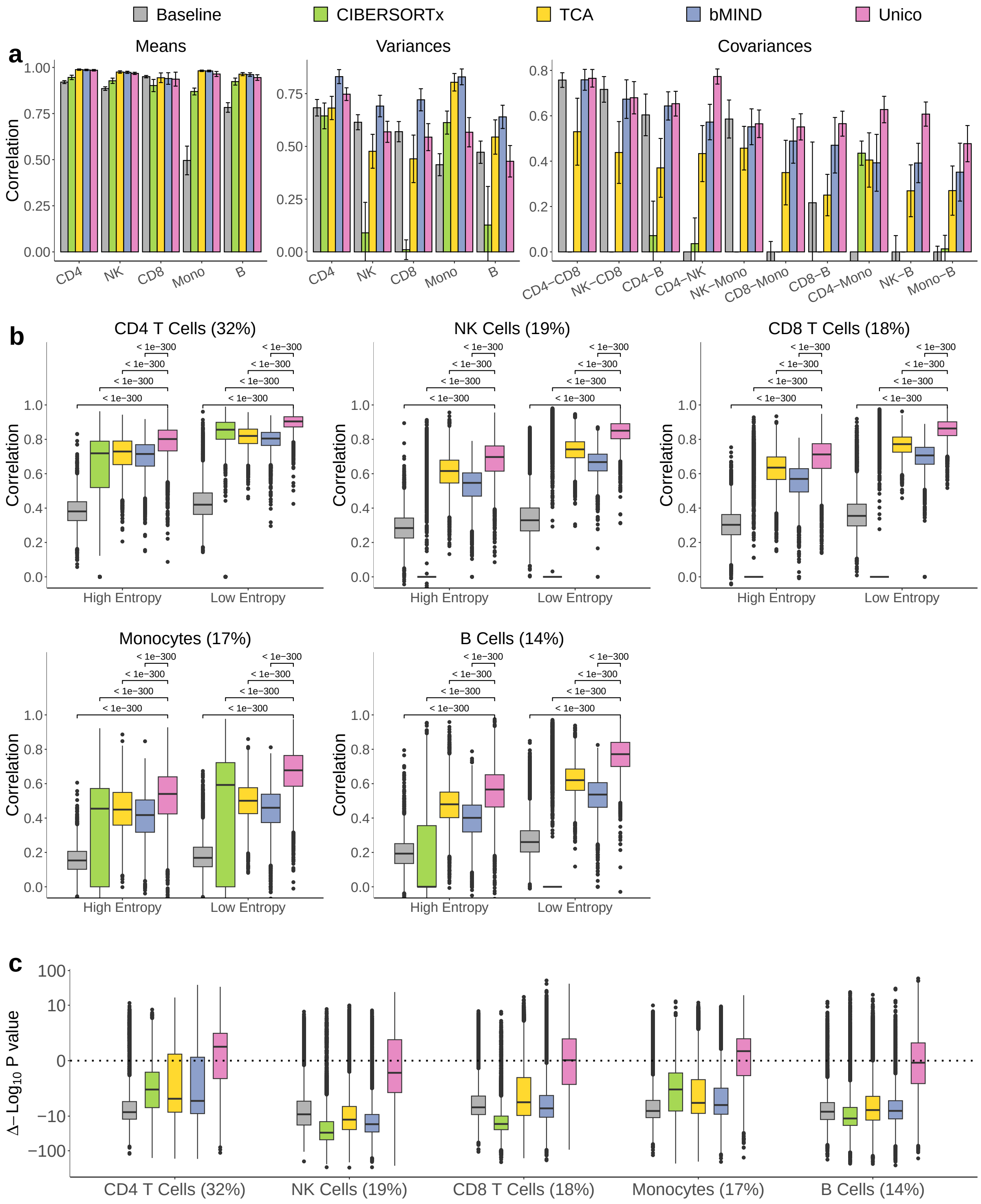

Figure S4: Evaluation of deconvolution methods on RNA pseudo-bulk mixtures. (a-c) Same analyses as in Supplementary Figure S1, but using only 250 samples in pseudo-bulk mixtures from PBMC scRNAseq profiles of five cell types (600 genes in each set)

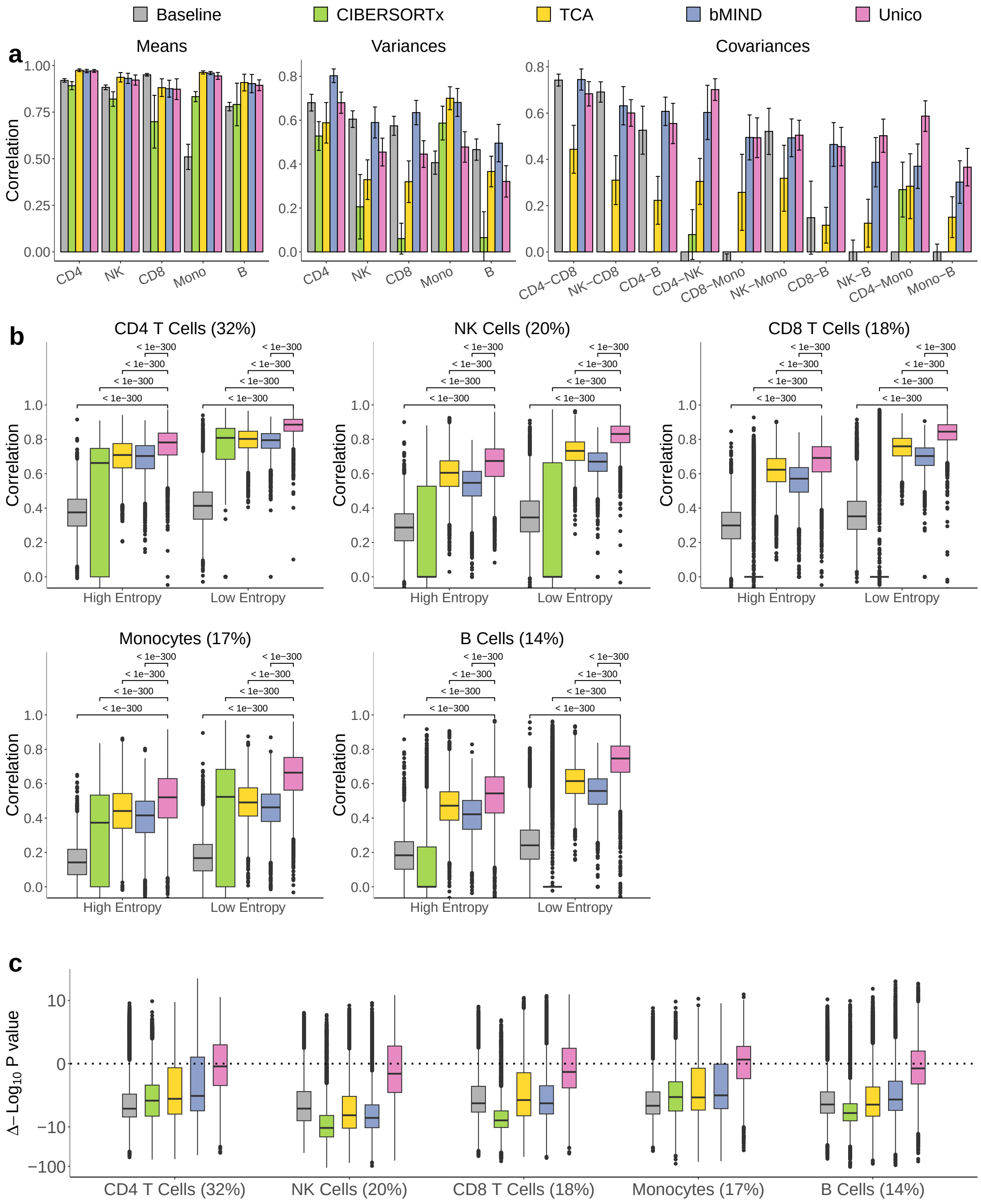

Figure S5: Evaluation of deconvolution methods on RNA pseudo-bulk mixtures. (a-c) Same analyses as in Supplementary Figure S1, but using only 100 samples in pseudo-bulk mixtures from PBMC scRNAseq profiles of five cell types (600 genes in each set)

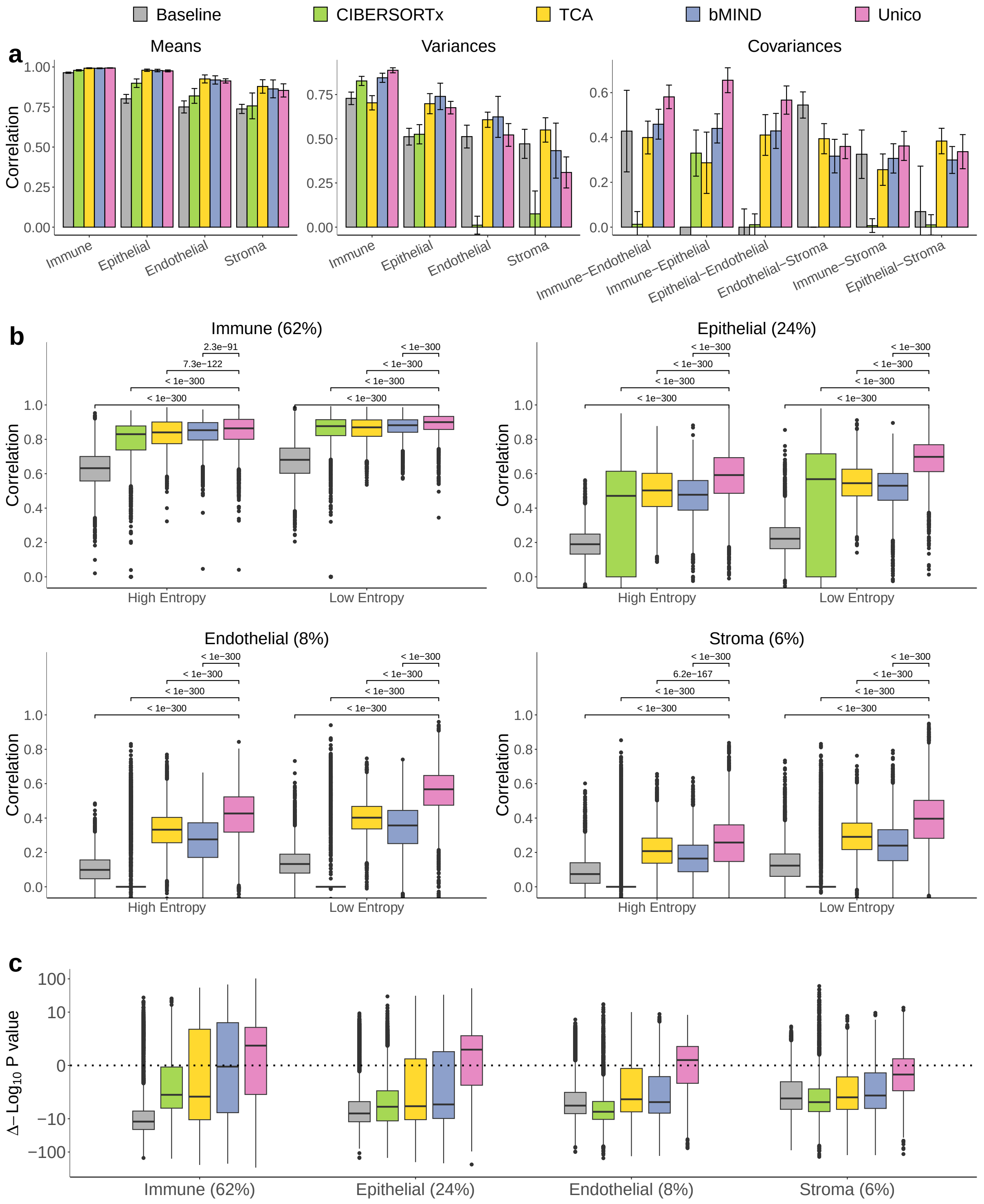

Figure S6: Evaluation of deconvolution methods on RNA pseudo-bulk mixtures. (a-c) Same analyses as in Supplementary Figure S1, but using only 250 samples in pseudo-bulk mixtures from lung scRNAseq profiles of four cell types (600 genes in each set)

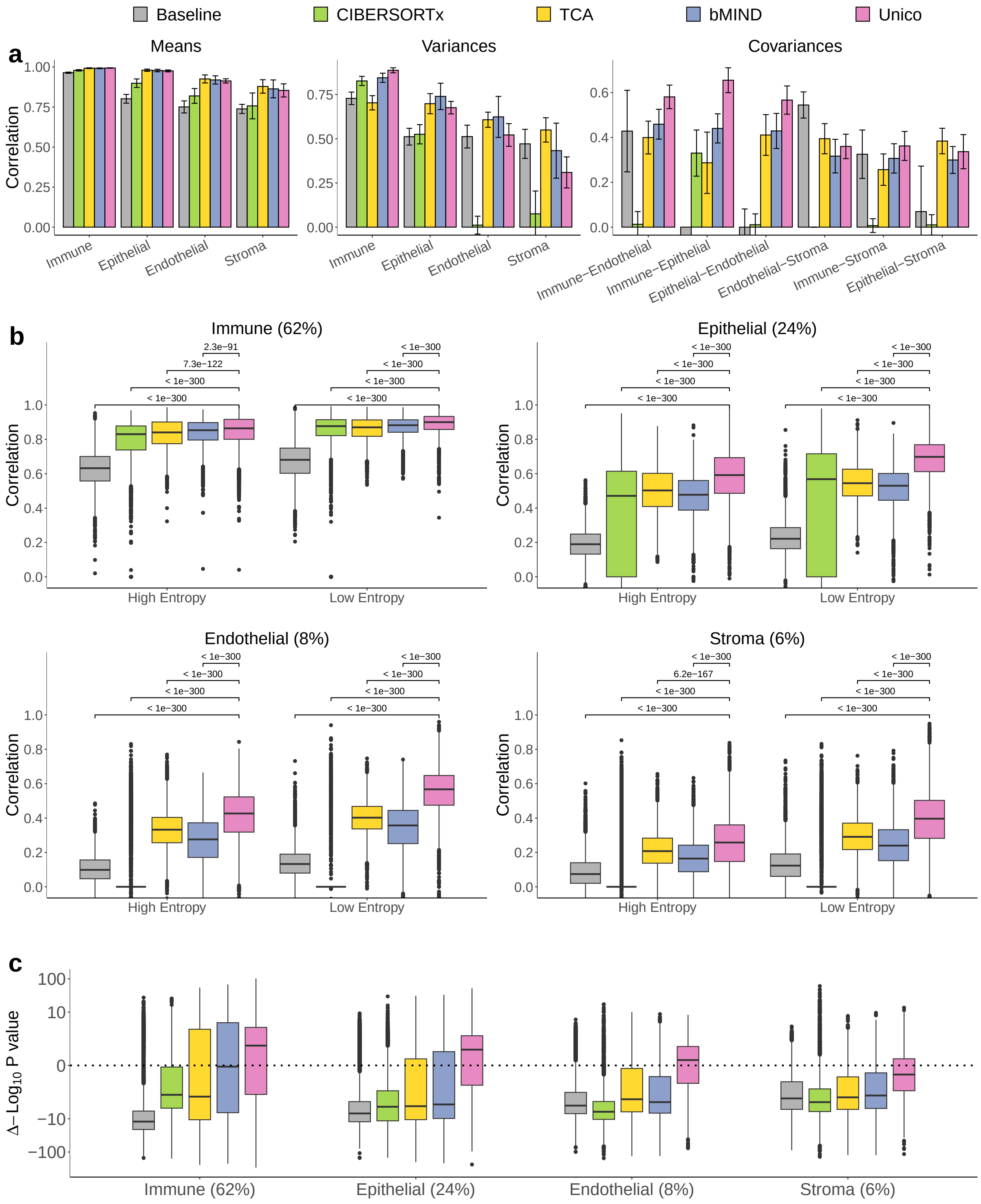

Figure S7: Evaluation of deconvolution methods on RNA pseudo-bulk mixtures. (a-c) Same analyses as in Supplementary Figure S1, but using only 100 samples in pseudo-bulk mixtures from lung scRNAseq profiles of four cell types (600 genes in each set)

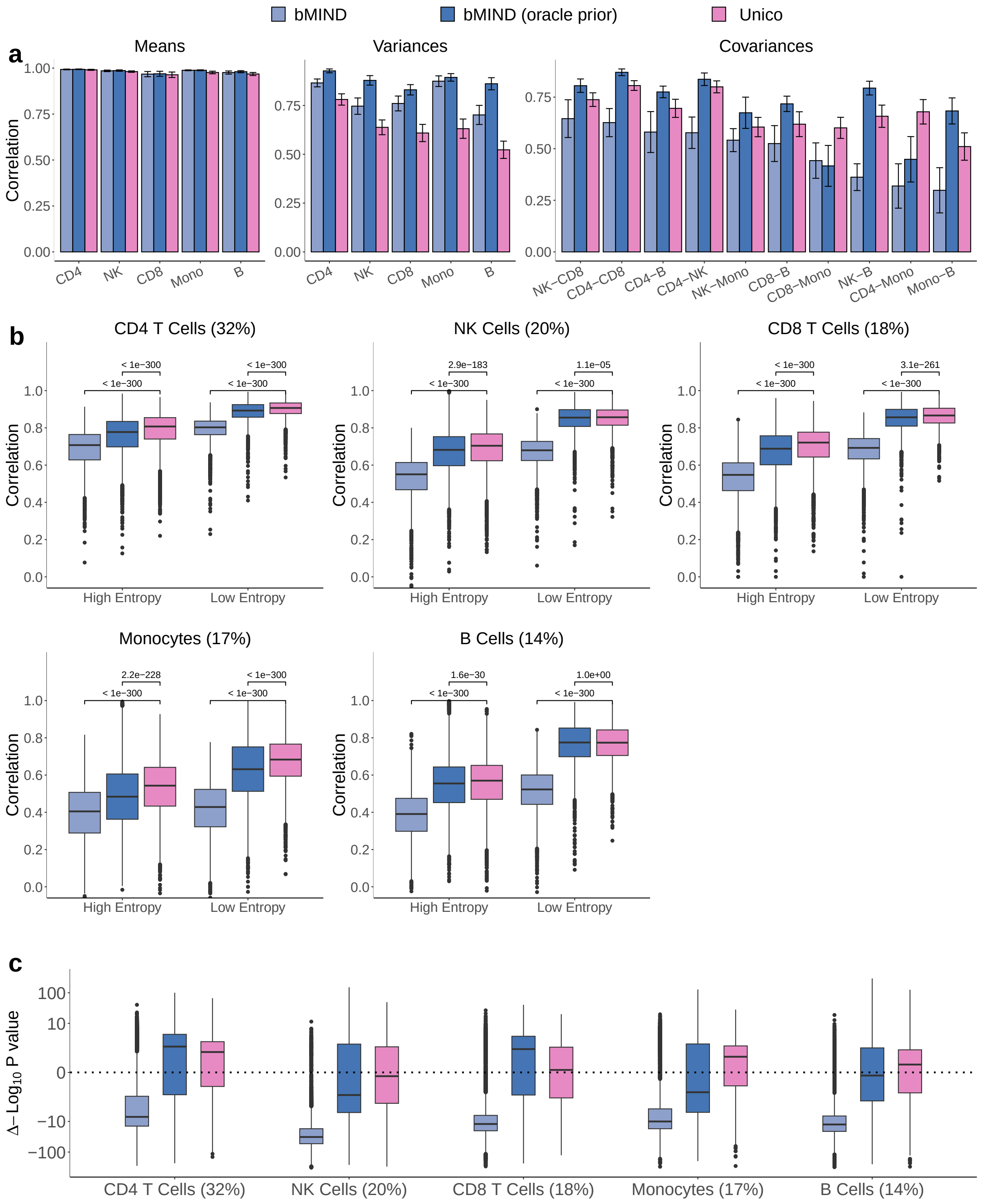

Figure S8: Evaluation of deconvolution methods on RNA pseudo-bulk mixtures. (a-c) Same analyses as in main Figure 2a-c, but also presenting bMIND in the presence of priors learned from the true cell-type levels of all samples in the PBMC scRNAseq dataset, denoted as “bMIND (oracle prior)” in dark blue, along with bMIND and Unico (500 samples and 600 genes in each set)

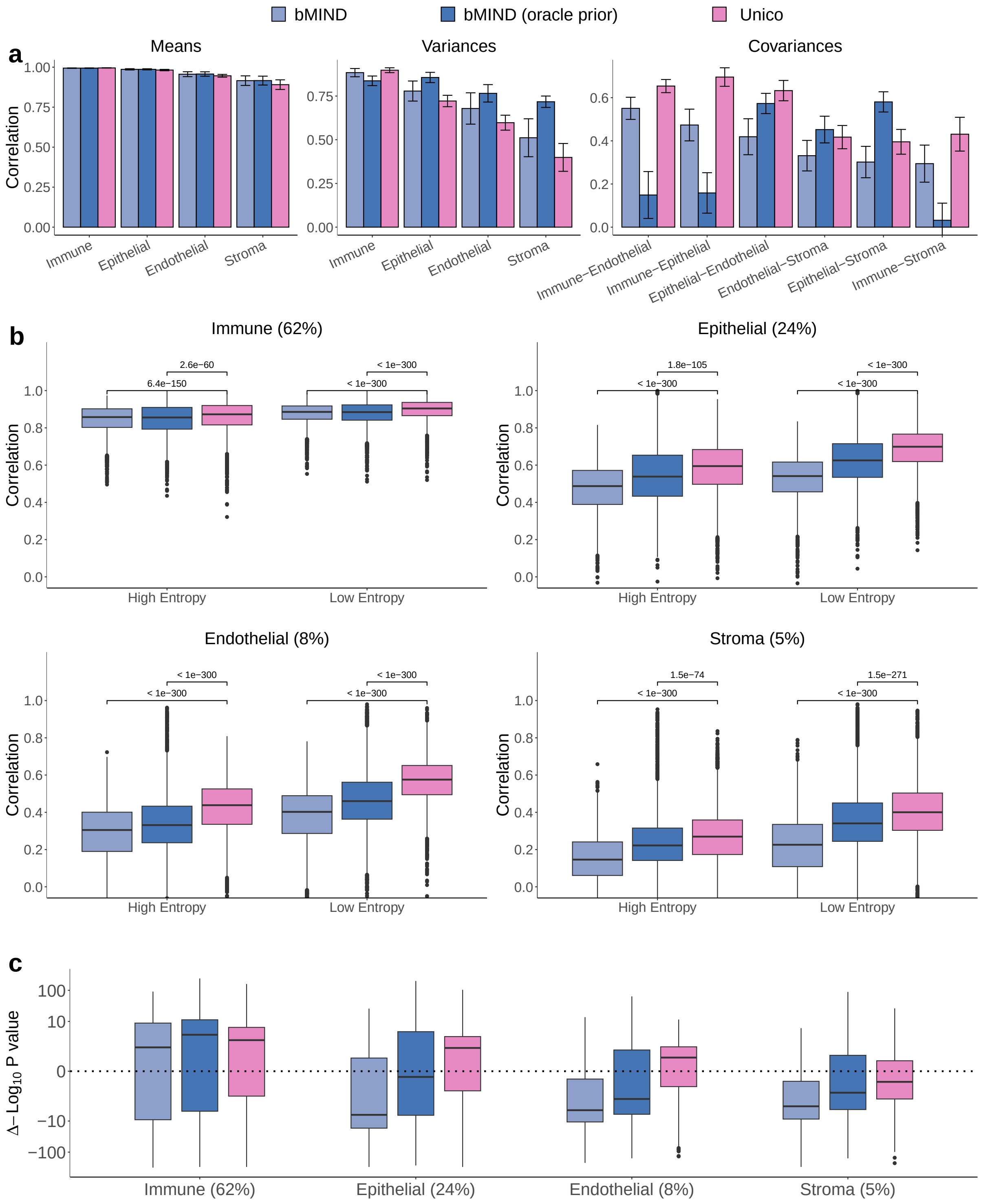

Figure S9: Evaluation of deconvolution methods on RNA pseudo-bulk mixtures. (a-c) The same analyses as in Supplementary Figure S1, but also presenting bMIND in the presence of priors learned from the true cell-type levels of all samples in the lung scRNAseq dataset, denoted as “bMIND (oracle prior)” in dark blue, along with bMIND and Unico (500 samples and 600 genes in each set)

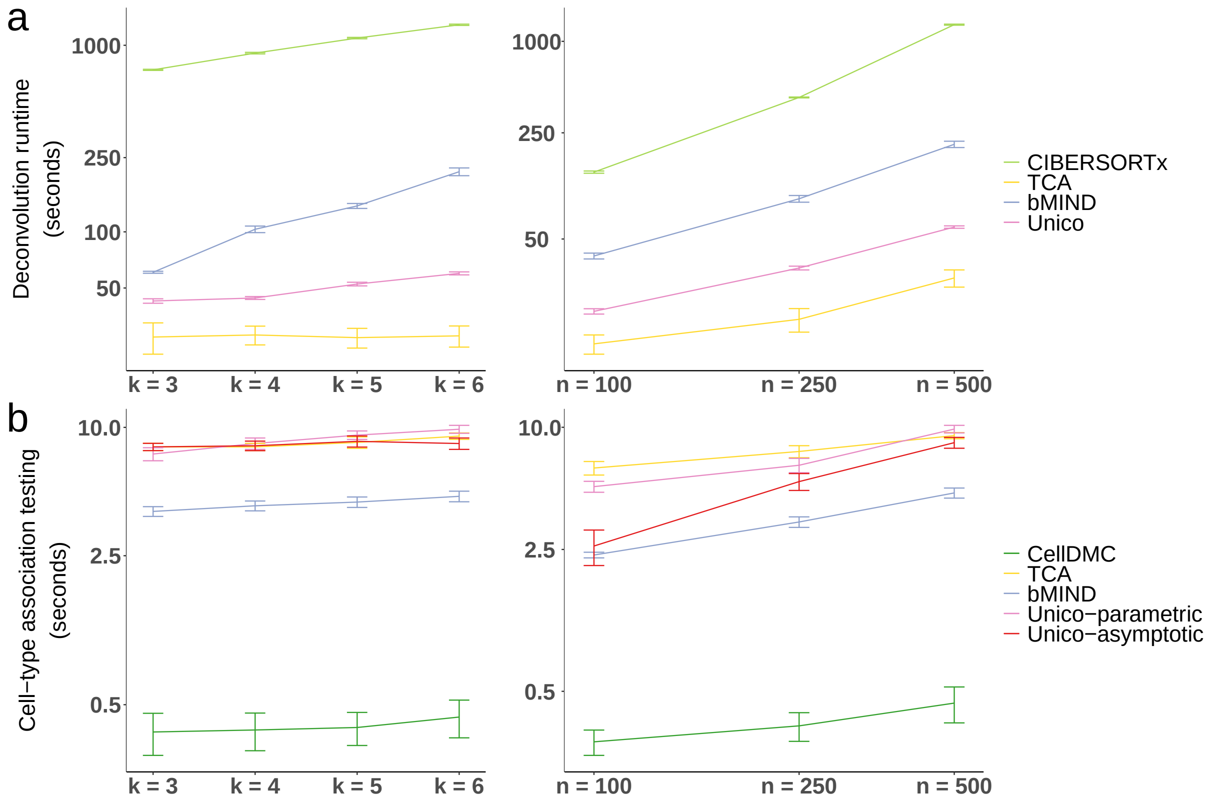

Figure S10: Runtime of the different deconvolution methods. (a) Evaluation of deconvolution runtime on bulk methylation profiles (500 samples, 1,000 CpGs) with a varying number of cell types. (b) Evaluation of deconvolution runtime on bulk methylation profiles (six cell types, 1,000 features) with a varying number of samples. (c)-(d) Similar to (a) and (b) only for cell-type level differential methylation testing. The varied number of cell types reflects different aggregations of cell types. All methods were executed on an ARM-based Apple M1 chip with 64G RAM and in parallel on 8 computational cores. In all plots, interval bars indicate the mean and one standard deviation of the runtime across 10 simulations (log-transformed scale). Samples were drawn at random from the Hannum et al. whole-blood methylation data [1] with $p_1 = 5$ cell-type level covariates and the top $p_2 = 10$ surrogates of technical variability as tissue-level covariates.

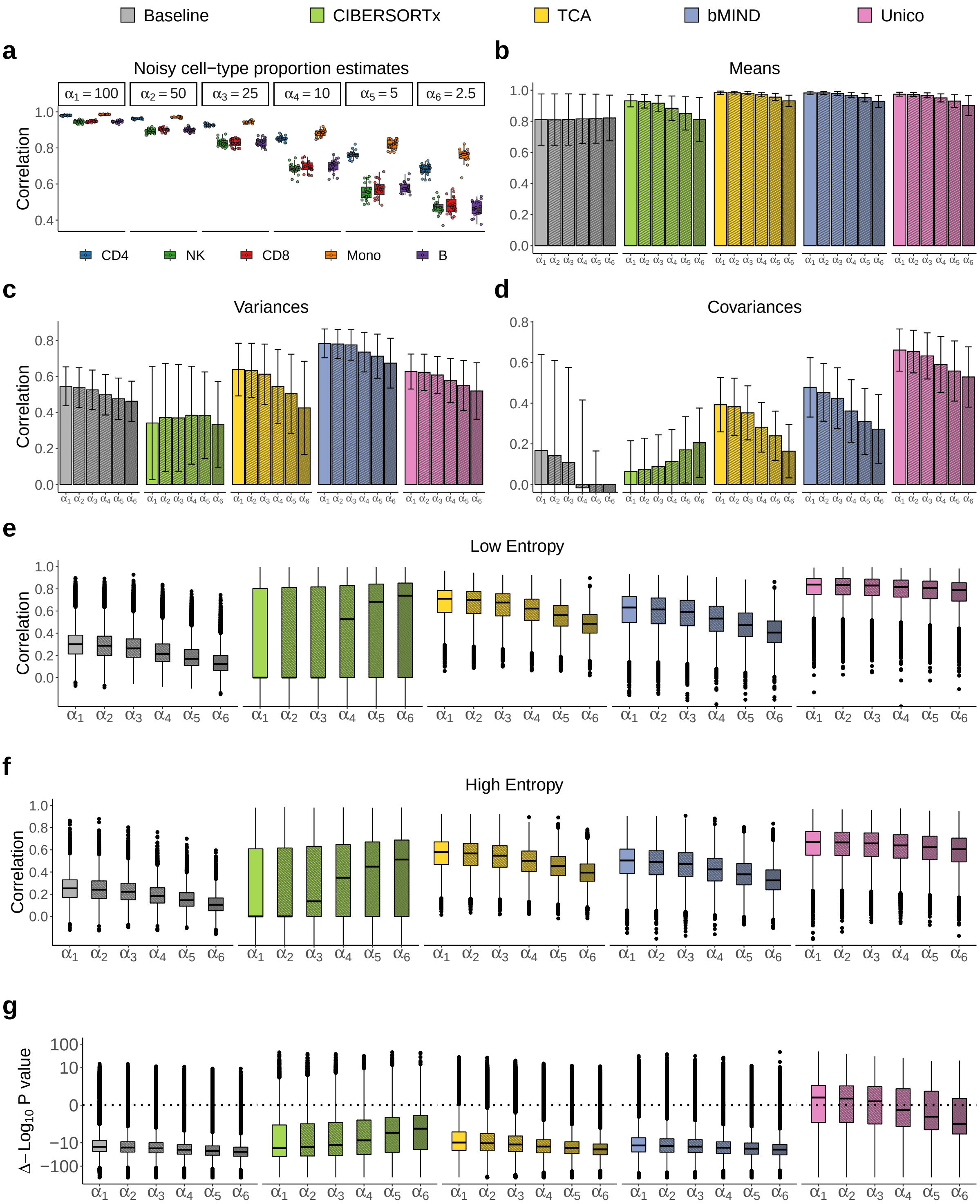
Baseline
CIBERSORTx
TCA
bMIND
Unico
a
Noisy cell−type proportion estimates
α1 = 100
α2 = 50
α3 = 25
α4 = 10
α5 = 5
α6 = 2.5
Correlation
CD4
NK
CD8
Mono
B
b
Means
c
Variances
Correlation
d
Covariances
e
Low Entropy
Correlation
f
High Entropy
Correlation
g
Δ−Log10 P value
α1
α2
α3
α4
α5
α6

Figure S11: Evaluation of deconvolution methods under varying levels of noise added to the cell-type proportions input. (a) Correlation between the ground truth cell-type proportions and the noisy version provided as input to all methods (lower $\alpha$ indicates more noise; Supplementary Methods). (b-d) Correlation between single-cell-based estimates of population-level cell-type moments and those based on deconvolution across 20 sets of pseudo-bulk mixtures from PBMC scRNAseq profiles (500 samples and 600 genes in each set). (e-f) Evaluation of the concordance between the known cell-type profiles and the deconvolution estimates. Boxplots reflect the distribution of linear correlation across all 5 cell types and all genes in the low entropy (e) and high entropy (f) set across the same 20 simulations in (b-d). (g) Assessing deconvolution methods for their information that cannot be explained by pseudo bulk expression. Boxplots reflect the distribution across all cell types and genes from the same data in (b-f) of $\Delta \log_{10}$(p-value), the difference between the log-scaled p-values of the effects of the pseudo bulk expression and deconvolution estimates (higher is better; Methods). All barplots and error bars in the figure represent means and one standard deviation errors; negative correlations were truncated for visualization purposes.

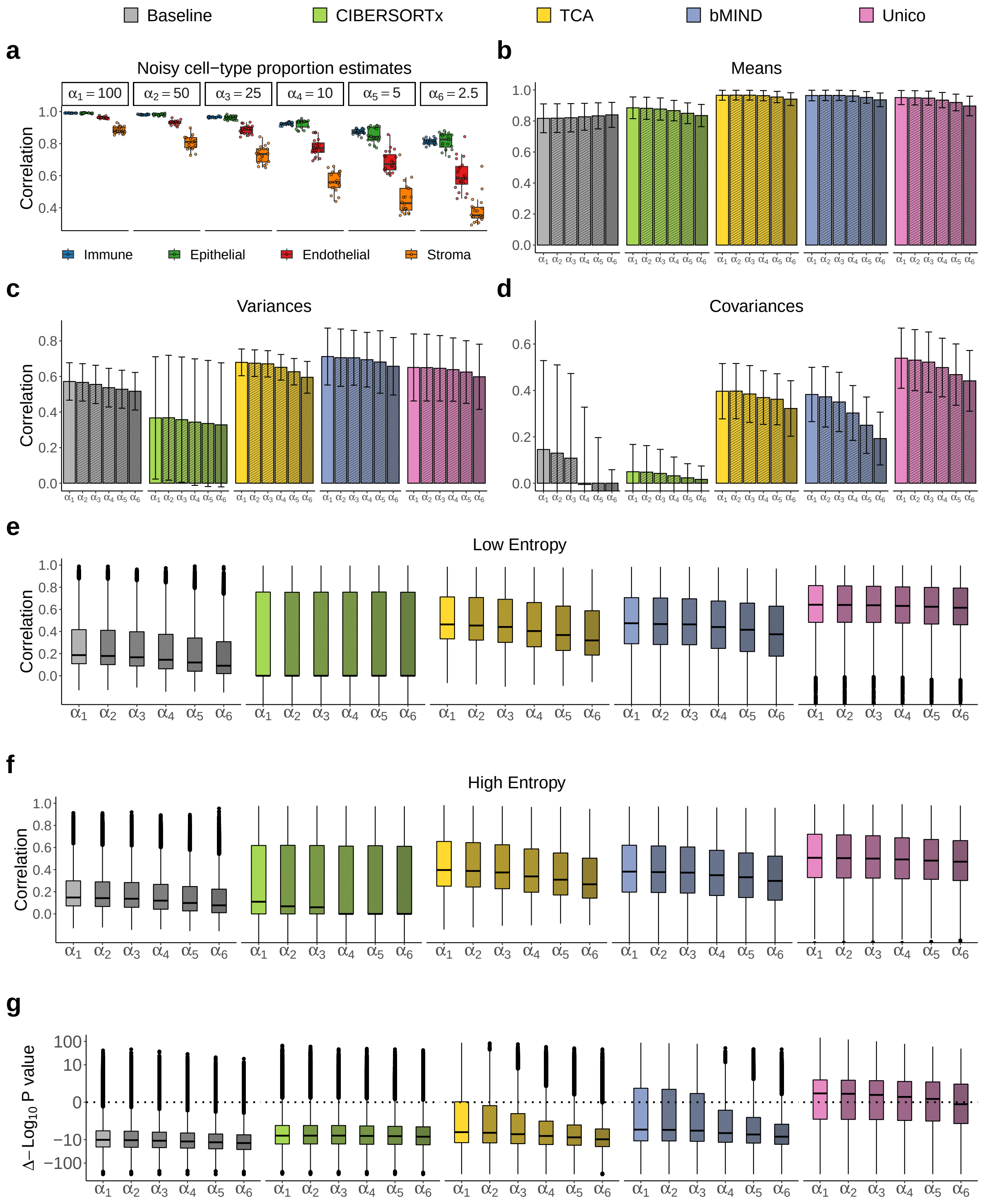

Figure S12: Evaluation of deconvolution methods under varying levels of noise added to the cell-type proportions input. Same analyses as in Supplementary Figure S11, but on pseudo-bulk mixtures from lung scRNAseq profiles of four cell types

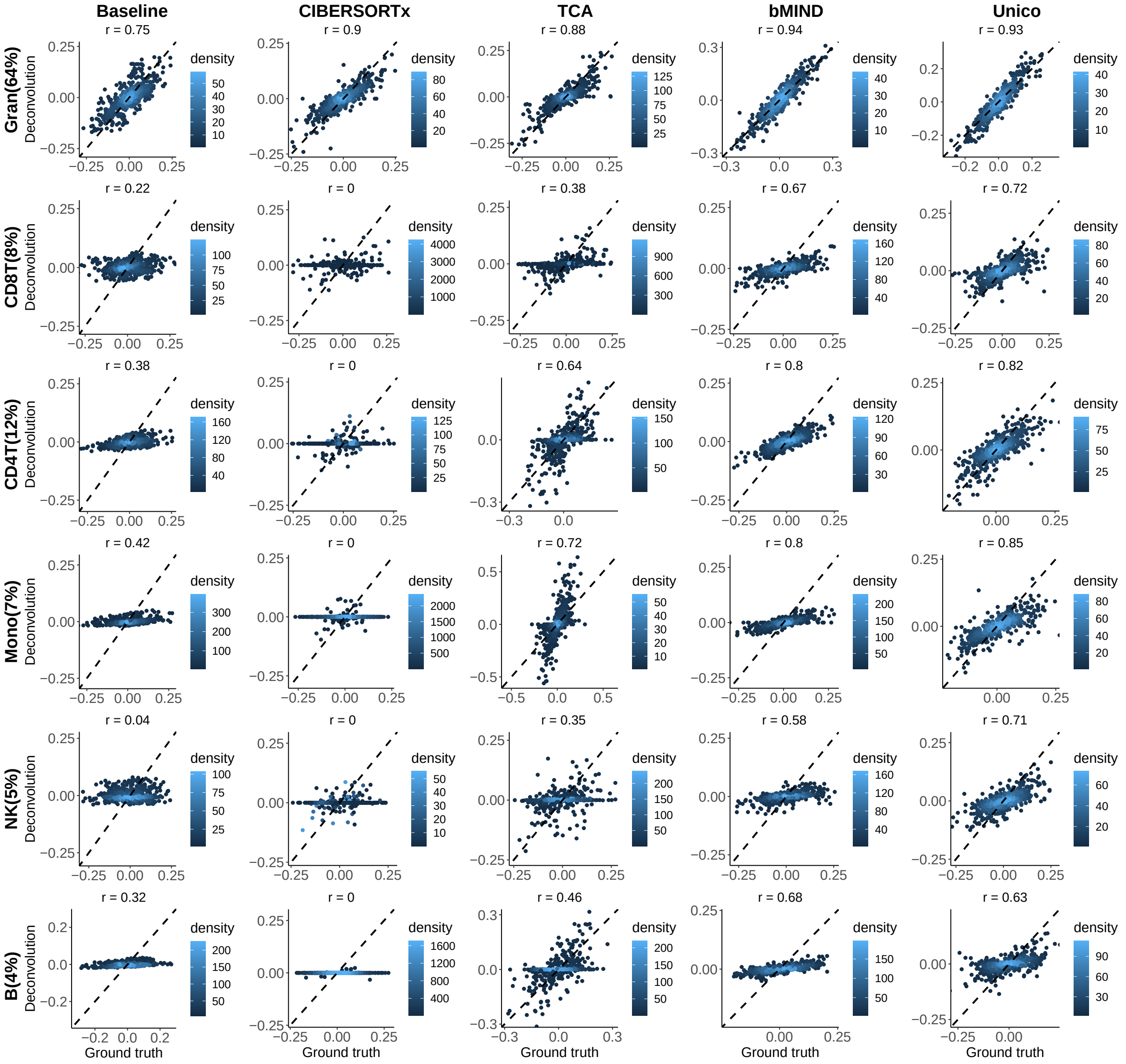

Figure S13: Deconvolution of the low-entropy CpGs in the set of 10,000 most variable CpGs in the Reinius whole-blood DNA methylation data. Presented are experimentally measured cell-type level methylation for the whole-blood samples (values pooled across samples and CpGs per cell type; "Ground truth") and the deconvolution estimates of the different methods ("Deconvolution").

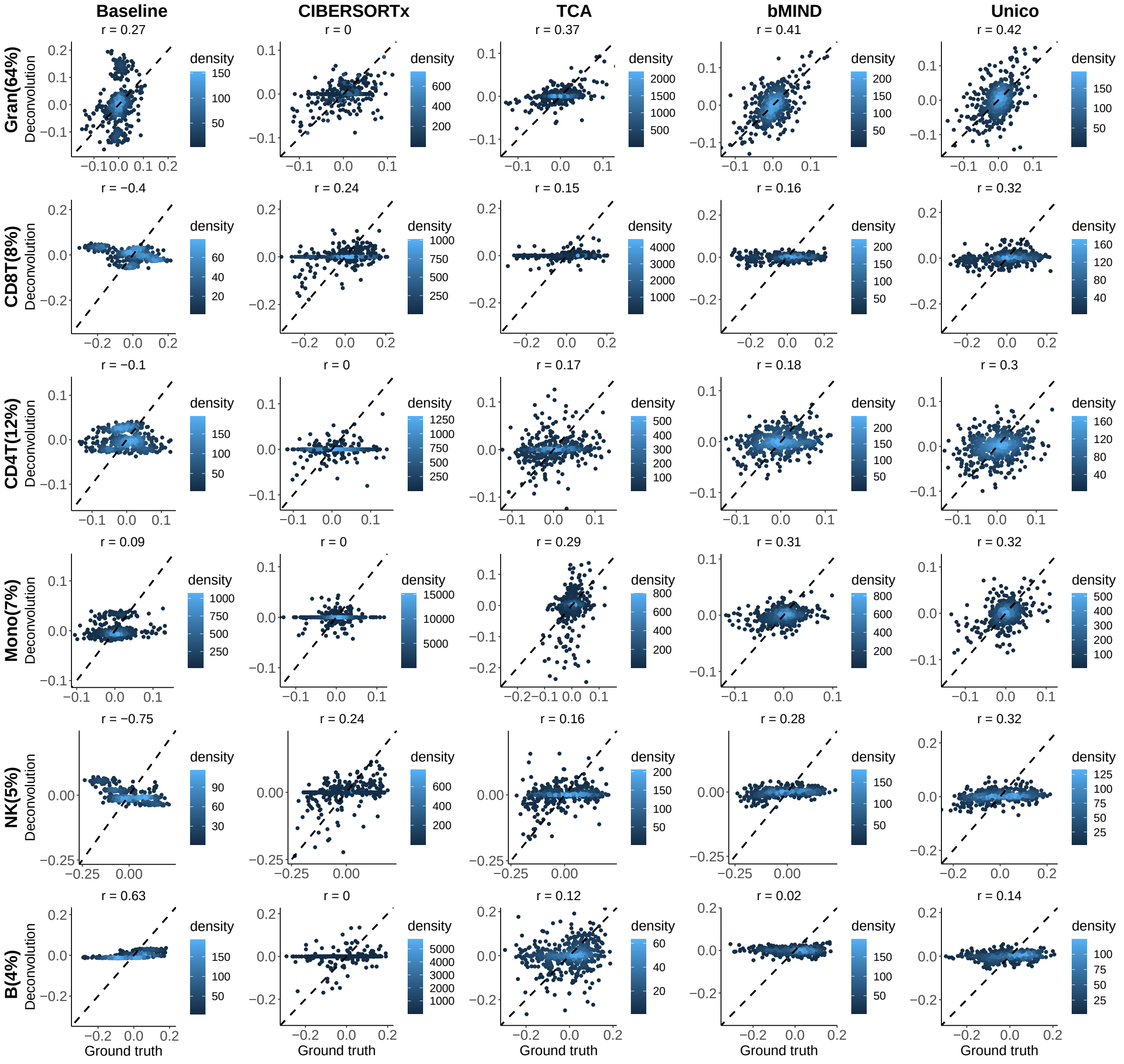

Figure S14: Deconvolution of the high-entropy CpGs in the set of 10,000 most variable CpGs in the Reinius whole-blood DNA methylation data. Presented are experimentally measured cell-type level methylation for the whole-blood samples (values pooled across samples and CpGs per cell type; "Ground truth") and the deconvolution estimates of the different methods ("Deconvolution").

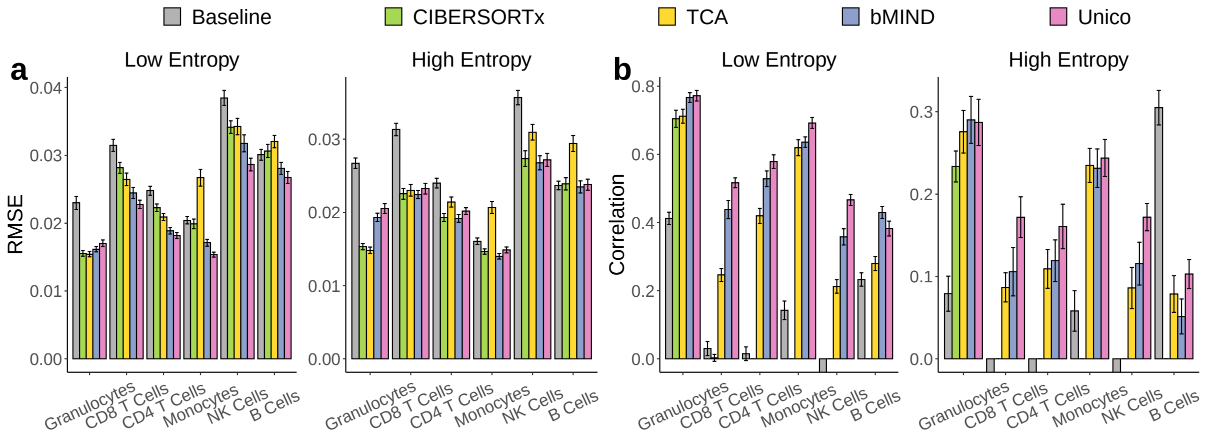

Figure S15: Evaluation of deconvolution methods in the set of 10,000 randomly selected CpGs in the Reinius whole-blood DNA methylation data. (a-b) Evaluation in terms of RMSE and correlation between estimates and experimentally validated cell-type level methylation across 20 random sets of 1,000 randomly selected CpGs. Barplots and error bars in the figure represent means and one standard deviation errors; negative correlations were truncated for visualization purposes.

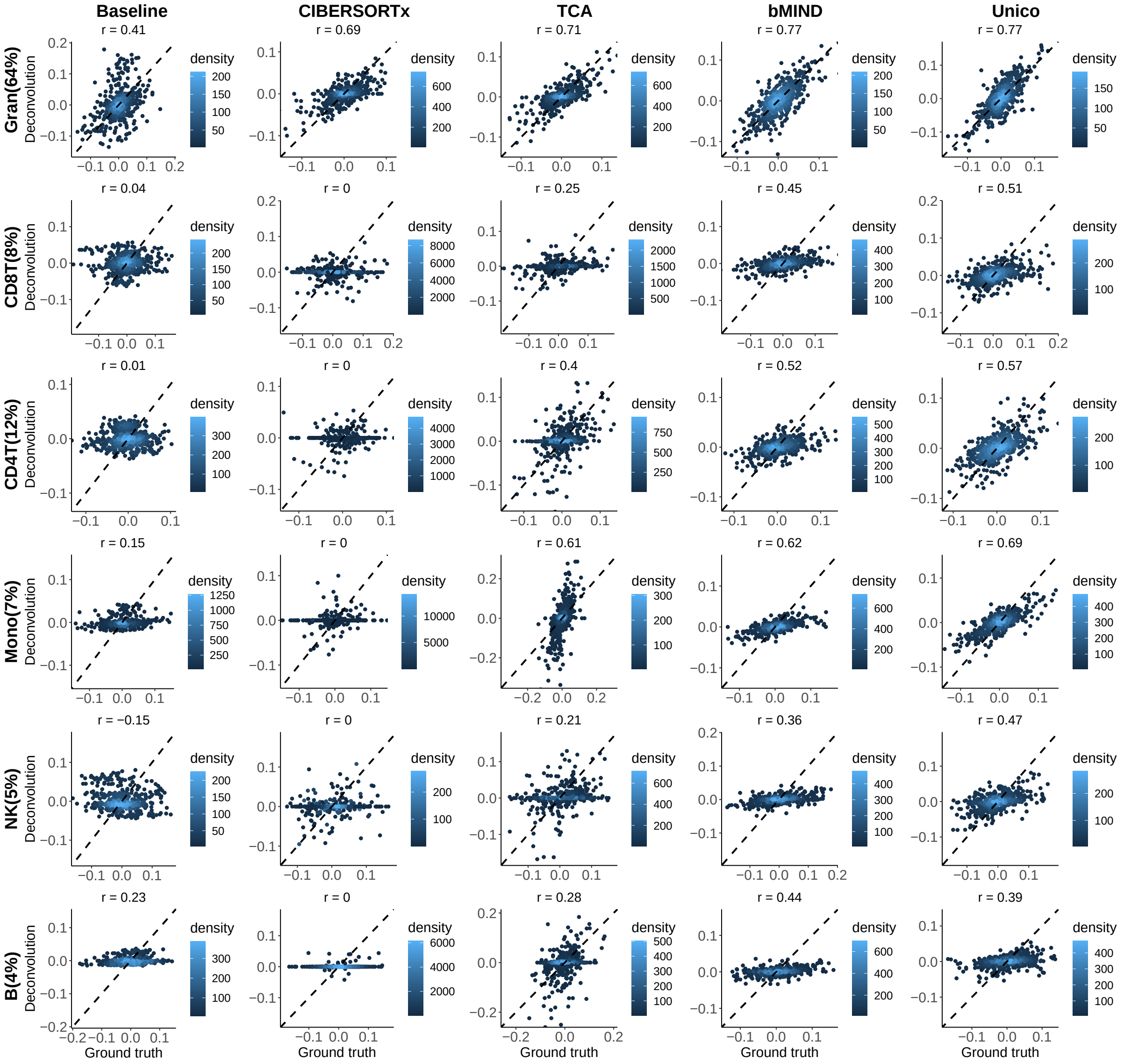

Figure S16: Deconvolution of the low-entropy CpGs in the set of 10,000 randomly selected CpGs in the Reinius whole-blood DNA methylation data. Presented are experimentally measured cell-type level methylation for the whole-blood samples (values pooled across samples and CpGs per cell type; "Ground truth") and the deconvolution estimates of the different methods ("Deconvolution").

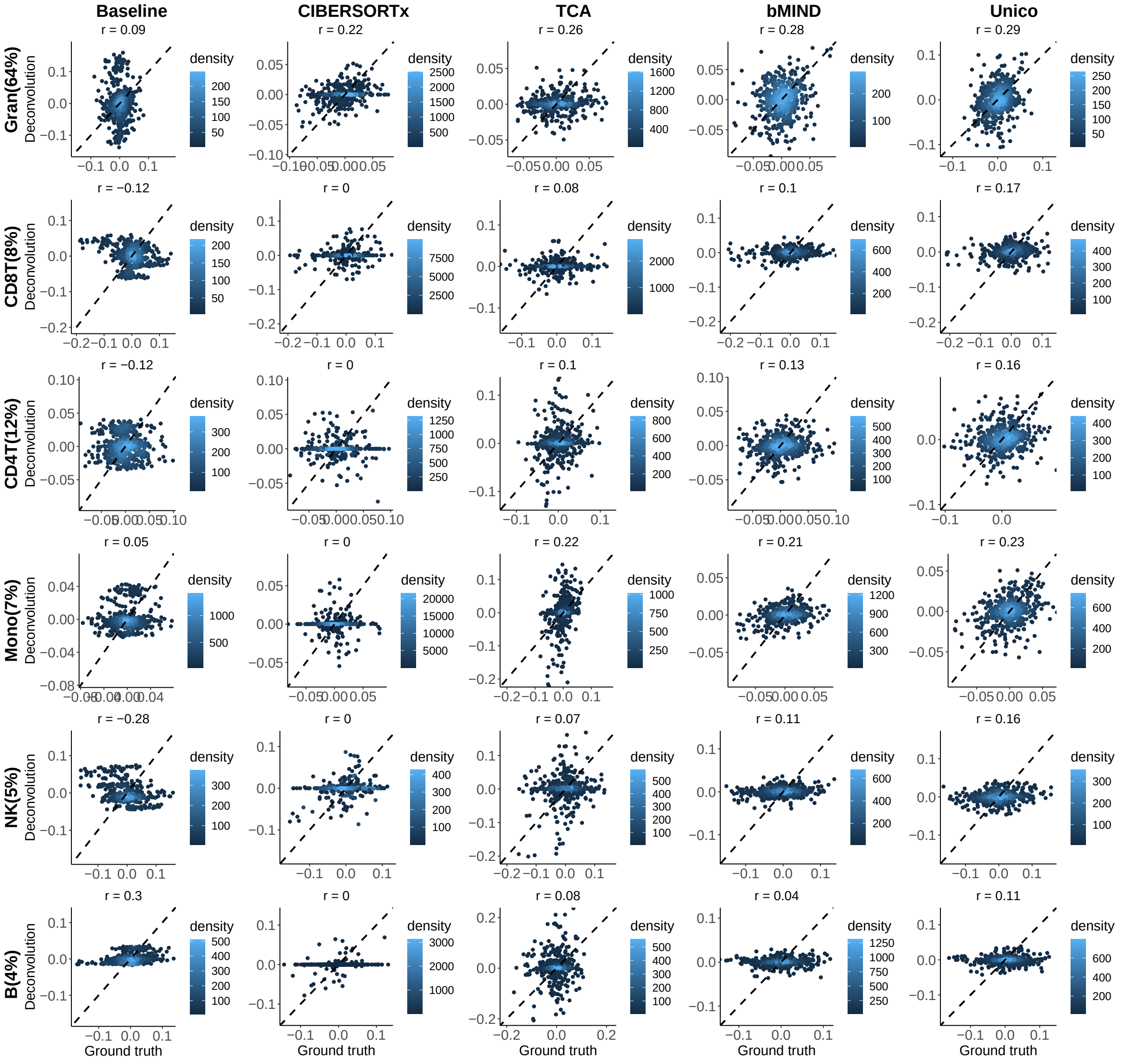

Figure S17: Deconvolution of the high-entropy CpGs in the set of 10,000 randomly selected CpGs in the Reinius whole-blood DNA methylation data. Presented are experimentally measured cell-type level methylation for the whole-blood samples (values pooled across samples and CpGs per cell type; "Ground truth") and the deconvolution estimates of the different methods ("Deconvolution").

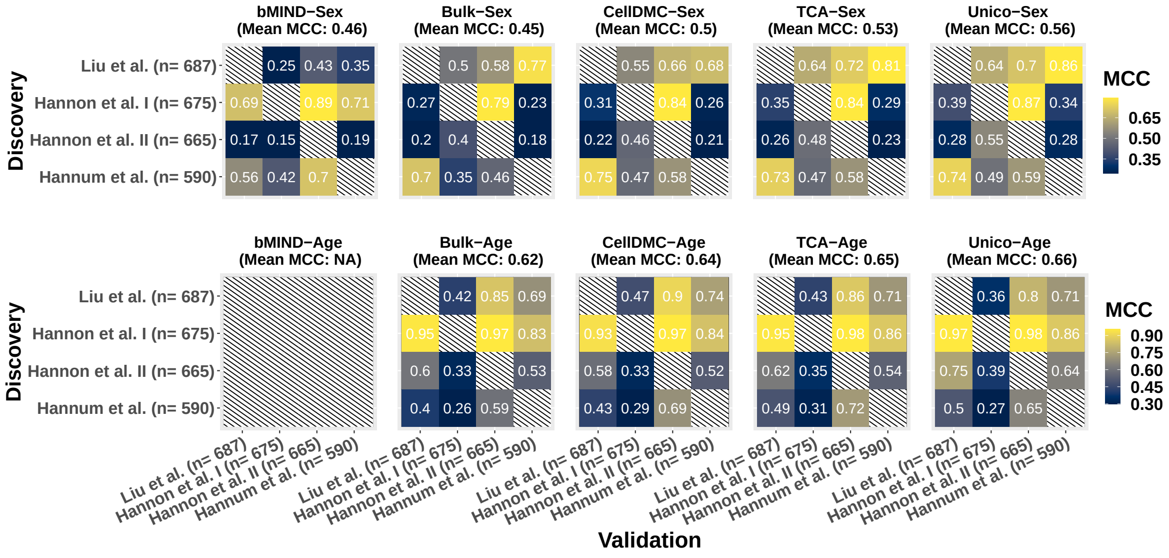

Figure S18: Consistency in calling tissue-level differential methylation with sex and age across four independent whole-blood DNA methylation datasets. Color gradients represent the Matthews correlation coefficient (MCC) for every possible pairing of two datasets as discovery and validation (Methods). Since bMIND was designed for binary conditions only, it was not evaluated in the age analysis

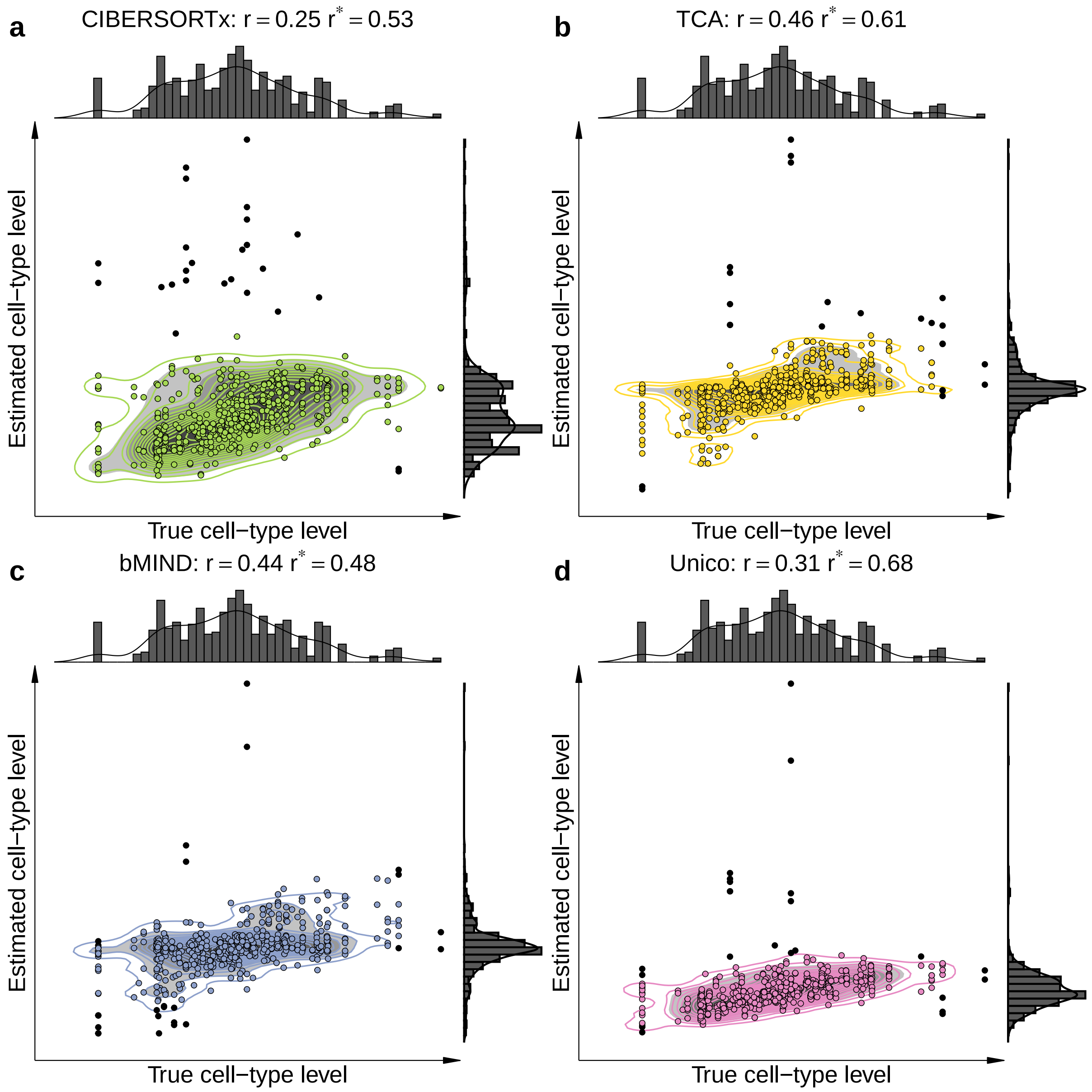

Figure S19: Example of the potential discrepancy between robust and non-robust correlation across different methods (a-d). Scatter plots show the concordance between the ground truth CD4 expression levels and their deconvolution-derived estimates for the gene SLC19A1 from the experiment of deconvolving RNA pseudo-bulk mixtures. Black-filled circles indicate 5% of the samples falling outside a 95% confidence ellipsoid (represented by contour lines) and are thus considered outliers. $r^*$ and $r$ indicate the sample linear correlation calculated with and without outliers removed, respectively.

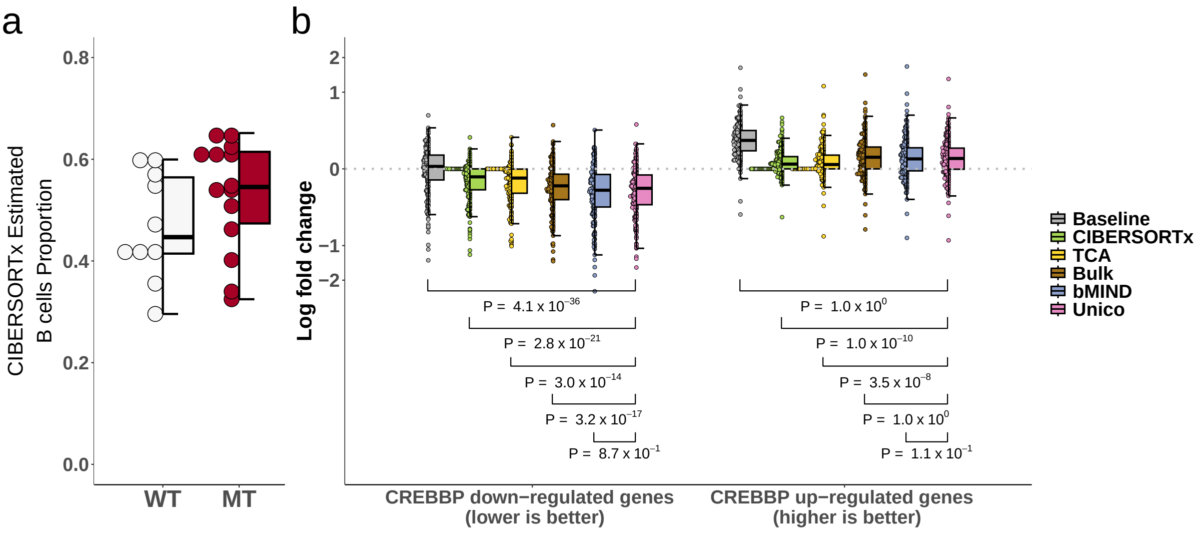

Figure S20: Deconvolution of bulk FL tumor samples. (a) B cell composition estimates from FL tumor samples with (MT; n=14) and without (WT; n=10) CREBBP mutation. (b) Deconvolution of bulk FL tumor samples for evaluating genes that were previously reported as differentially expressed with CREBBP mutation in B cells of FL tumors. Presented are the log (basis 2) fold change across 219 down-regulated and 275 up-regulated genes, evaluated on B cell expression estimated by deconvolving the bulk FL samples; pairwise method comparisons against Unico were assessed using a one-sided paired Wilcoxon test.

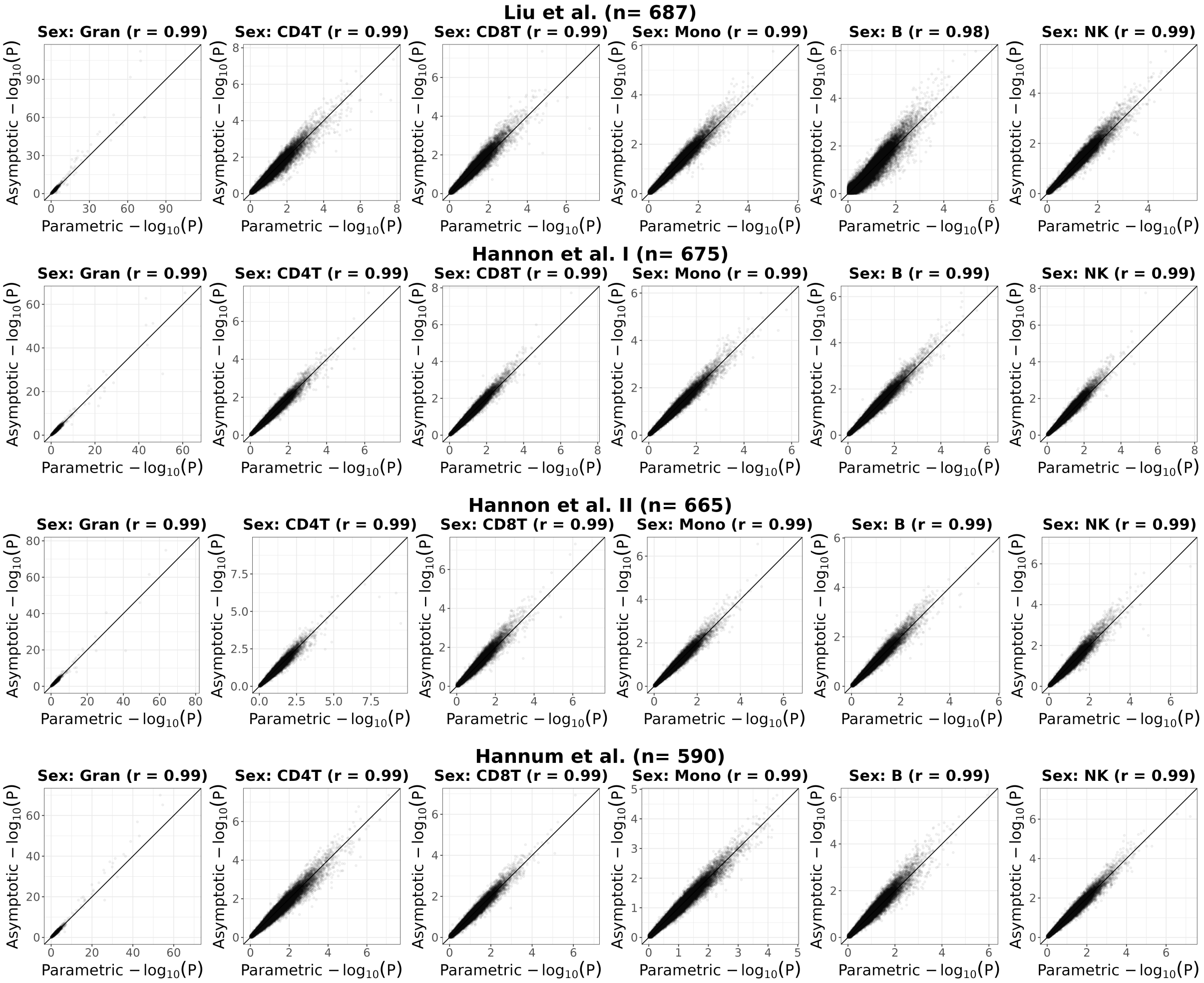

Figure S21: Evaluation of Unico's asymptotically-derived p-values under non-parametric testing for cell-type level differential methylation with sex in four whole-blood datasets. Presented are scatter plots showing log-transformed p-values under the assumption that methylation levels are normally distributed ("Parametric") versus the corresponding log-transformed p-values of a non-parametric test ("Asymptotic").

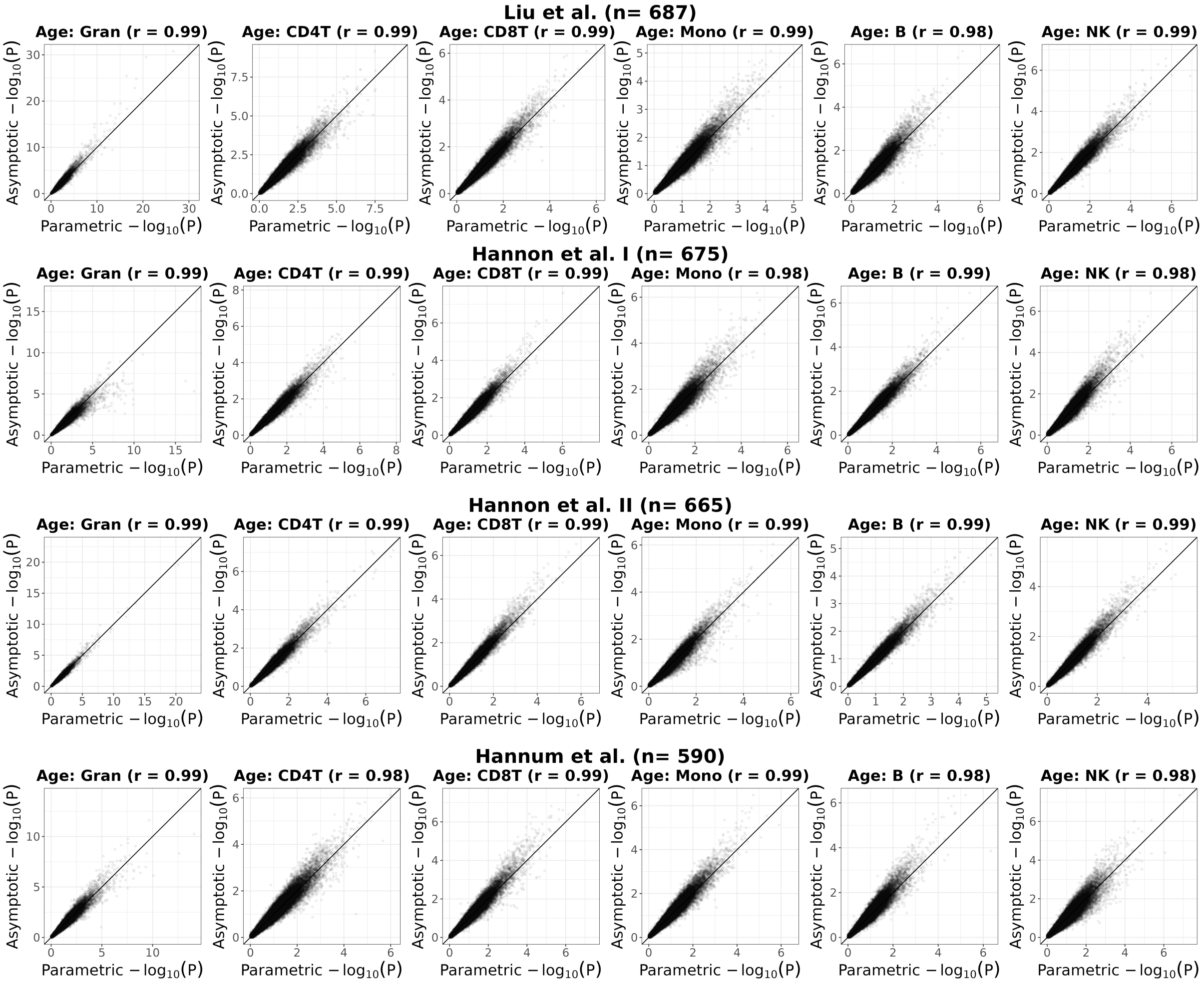

Figure S22: Evaluation of Unico's asymptotically-derived p-values under non-parametric testing for cell-type level differential methylation with age in four whole-blood datasets. Presented are scatter plots showing log-transformed p-values under the assumption that methylation levels are normally distributed ("Parametric") versus the corresponding log-transformed p-values of a non-parametric test ("Asymptotic").

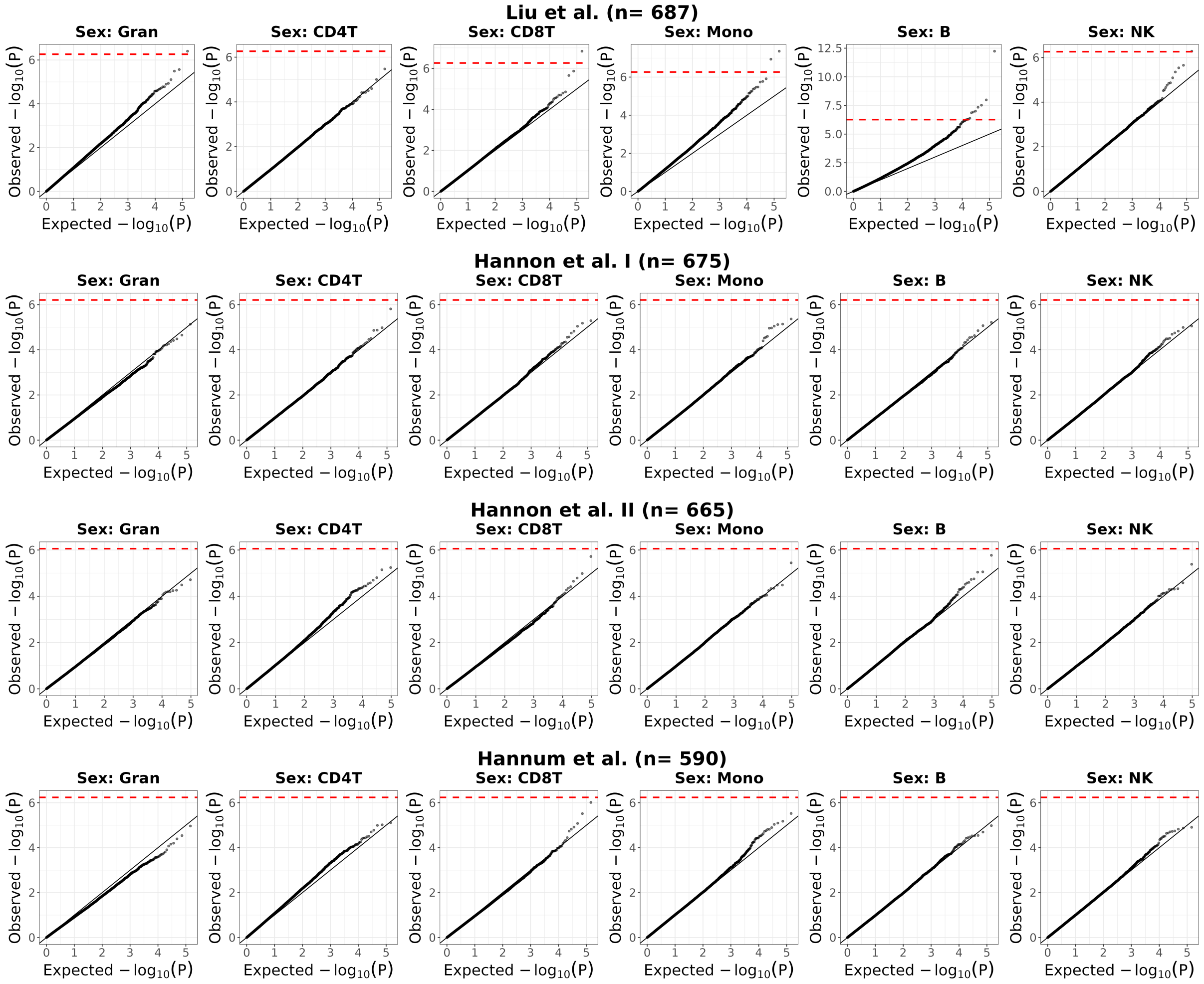

Figure S23: Evaluation of the null distribution of Unico's asymptotically derived p-values under non-parametric testing for cell-type level differential methylation with sex in four whole-blood datasets. Presented are quantile-quantile plots with log-transformed expected p-values versus the observed p-values under permutations of the condition (i.e., under the null). Red horizontal dashed lines indicate the Bonferroni-corrected threshold, adjusting for the number of CpGs and the number of cell-types under test.

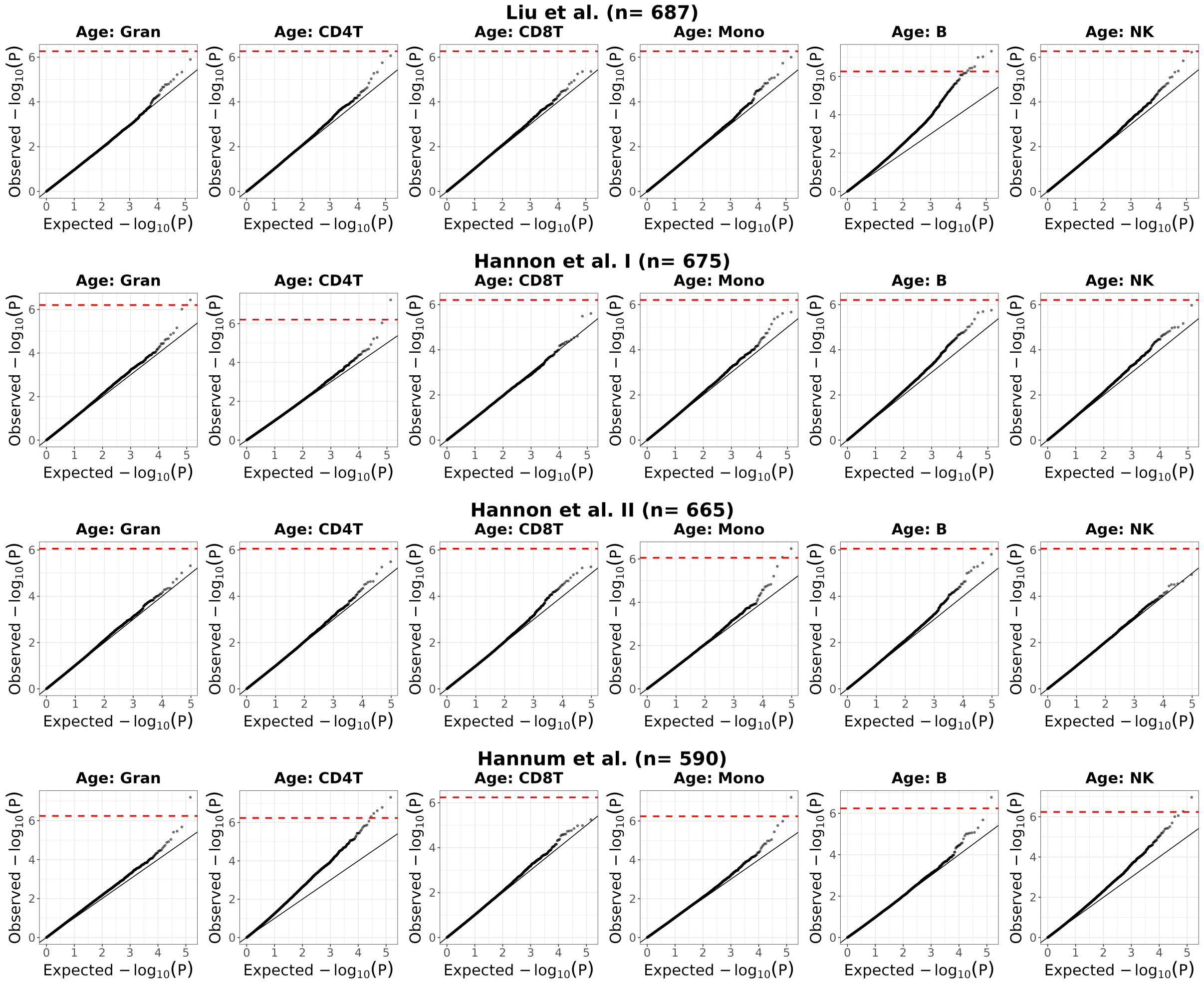

Figure S24: Evaluation of the null distribution of Unico's asymptotically derived p-values under non-parametric testing for cell-type level differential methylation with age in four whole-blood datasets. Presented are quantile-quantile plots with log-transformed expected p-values versus the observed p-values under permutations of the condition (i.e., under the null). Red horizontal dashed lines indicate the Bonferroni-corrected threshold, adjusting for the number of CpGs and the number of cell types under test.

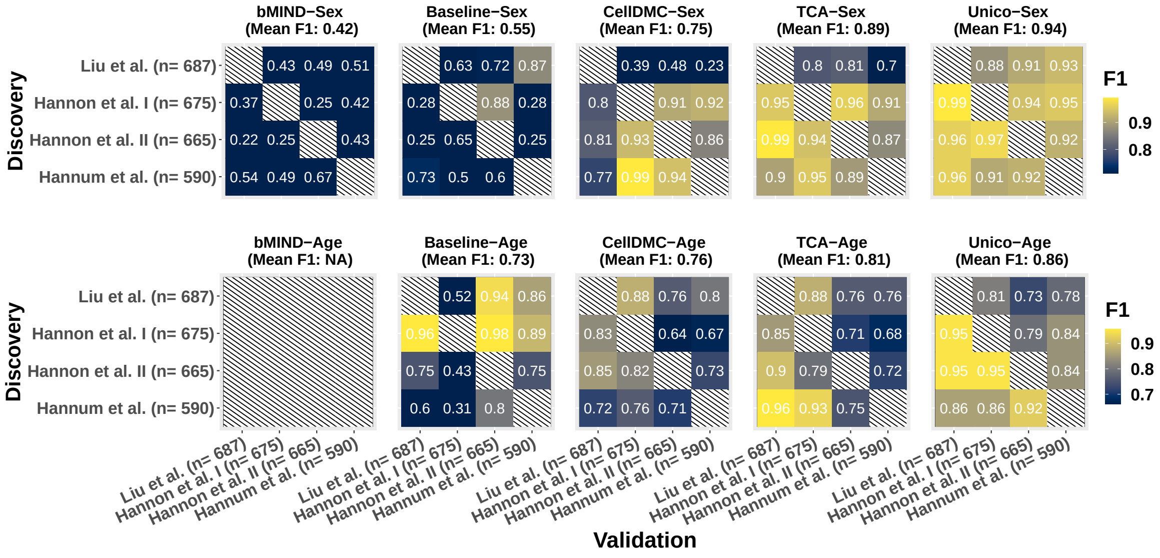

Figure S25: Consistency in calling cell-type level differential methylation with sex and age across four independent whole-blood DNA methylation datasets. Color gradients represent the F1 score for every possible pairing of two datasets as discovery and validation (Methods). Since bMIND was designed for binary conditions only, it was not evaluated in the age analysis

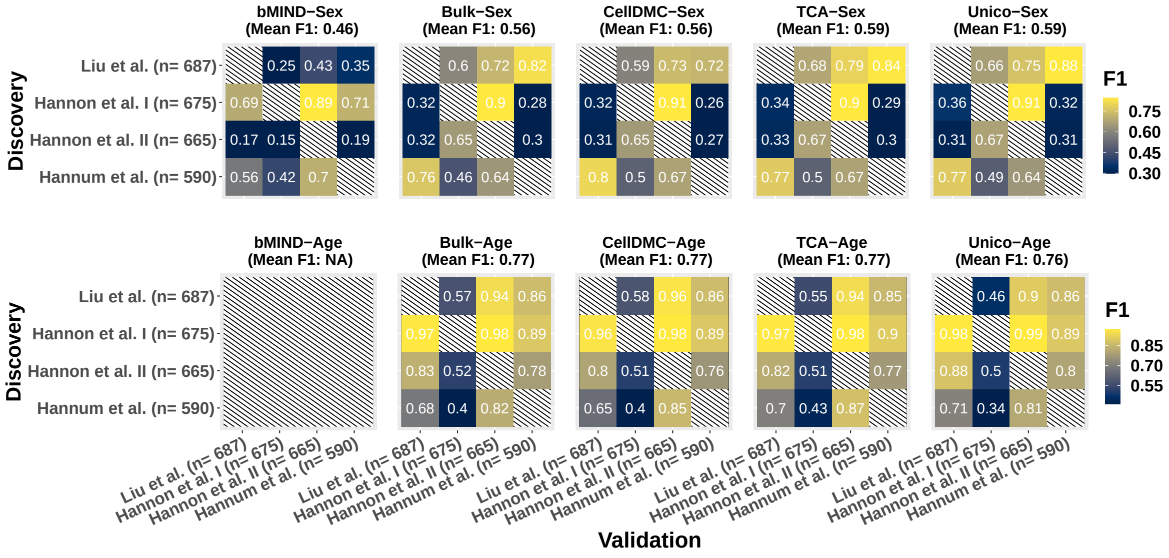

Figure S26: Consistency in calling tissue-level differential methylation with sex and age across four independent whole-blood DNA methylation datasets. Color gradients represent the F1 score for every possible pairing of two datasets as discovery and validation (Methods). Since bMIND was designed for binary conditions only, it was not evaluated in the age analysis

### S2 Supplementary Methods

#### S2.1 Unico: uniform cross-omics deconvolution

##### S2.1.1 Unico: distribution-free deconvolution incorporating cell-type covariance Modeling

Let $X_{ij}$ be the (tissue-level) bulk level of sample $i \in \{1...,n\}$ in feature $j \in \{1...,m\}$, and let $Z_{ij} = (Z_{ij1},...,Z_{ijk})$ be the levels of sample $i$ in feature $j$ and each cell type $h \in \{1...,k\}$, the full Unico model assumes:

$$X_{ij} = w_i^T Z_{ij} + (c_i^{(2)})^T \beta_j + e_{ij} \quad (1)$$

$$E[e_{ij}] = 0, V[e_{ij}] = \tau_j^2 \quad (2)$$

$$Z_{ijh} = \mu_{jh} + (c_i^{(1)})^T \gamma_{jh} + \epsilon_{ijh} \quad (3)$$

$$E[\epsilon_{ijh}] = 0, V[\epsilon_{ijh}] = \sigma_{jh}^2 \quad (4)$$

$$\sigma_{jh,jq} \equiv \text{Cov}[Z_{ijh}, Z_{ijq}], \sigma_{jh,jh} \equiv \sigma_{jh}^2 \quad (5)$$

where $w_i = (w_{i1},...,w_{ik})$ is a vector of sample-specific cell-type proportions of $k$ cell types that are assumed to compose the studied tissue, $e_{ij}$ is an i.i.d. component of variation that reflects measurement noise, $c_i^{(2)}$ is a $p_2$-length vector of known covariate values of sample $i$ that may demonstrate global effects (e.g., batch effects), and $\beta_j$ is a vector of the corresponding gene-specific fixed effect sizes. The parameters $\mu_{jh}$ represent mean levels, specific to gene $j$ and cell type $h$, $\epsilon_{ihj}$ is an i.i.d. noise term with mean zero and variance $\sigma_{jh}^2$ that may be specific to gene $j$ and cell type $h$, $c_i^{(1)}$ is a $p_1$-length vector of known covariate values of sample $i$ that may demonstrate cell-type-specific effects, and $\gamma_{jh}$ is a vector of the corresponding fixed effect sizes.

Equation (5) reflects our expectation that at least some genomic features will be different yet coordinated between different cell types. Particularly, transcriptional programs are known to persist through multiple differentiation steps, thus leading to differences in expression between cell lineages [2, 3]. Indeed, we observe that a smaller distance on the lineage differentiation tree corresponds to a higher degree of correlation. For example, using large single-cell data from peripheral blood mononuclear cells (PBMCs) (n=118 individuals) [4],

Among the top 10,000 most highly expressed genes in the single-cell PBMC data, we found that the median correlation between monocytes (myeloid lineage) and CD8 T cells (lymphoid lineage) is 0.25. In contrast, different lymphoid cell types tend to be more correlated: for example, the median correlation between CD4 T cells and CD8 T cells is 0.71. The median correlation between monocytes and CD4 T cells, B cells, and NK cells are 0.27, 0.30, and 0.37, respectively. The median correlation between CD4 T versus NK cells, CD4 T versus B cells, CD8 T versus NK cells, CD8 T versus B cells, and NK versus B cells are 0.48, 0.40, 0.46, 0.36, and 0.45, respectively.

###### S2.1.2 Unico in the context of previous deconvolution methods

The first method proposed in the space of deconvolution of transcriptomics is CIBERSORTx transcriptomics [5]. CIBERSORTx uses a heuristic approach based on the key idea that if a data point is used in multiple non-negative matrix factorization problems of different subsets of the data, then its underlying cell-type-specific signals can be approximated by a low-rank combination of the different decomposition products.

TCA is the first deconvolution method proposed for DNA methylation [6]. TCA employs a likelihood-based optimization that was specifically tailored for DNA methylation data, for which the normality assumption is arguably proper for the vast majority of CpGs if collected from methylation arrays. More concretely, the model follows Equations (1)-(4), while making the additional assumption that the i.i.d. component of variation $e_{ij}$ in Equation (2) is normally distributed and for a given feature $j$ and cell type $h$ the random variables $\{Z_{ijh}\}$ follow a normal distribution with no covariance structure assumed. Unico can therefore be viewed as a generalization of the TCA model, as the former does not make distributional assumptions and further models cell-type covariance. Furthermore, since standard decomposition was shown to be a degenerate case of TCA [7], Unico can also be viewed as a generalization of standard decomposition.

Similarly to TCA, the bMIND [8] model follows Equations (1)-(4), and in addition Equation (5), which models cell-type covariance, while making the additional assumption that the components of variation are normally distributed. However, unlike other deconvolution methods, bMIND is Bayesian and assumes the data is normally distributed. bMIND can incorporate priors for the means and covariances, which can be estimated from either independent single-cell data or the analyzed bulk data, and the model is learned using Markov chain Monte Carlo (MCMC) sampling for optimization.

Finally, we would like to note that CODEFACS [9] and MIND [10] were also proposed as deconvolution methods in the context of transcriptomics. CODEFACS is a heuristic method, which was neither open-sourced nor fully detailed by the authors, and MIND is a model-based method that requires multiple measurements from the same samples/individuals, which is beyond the scope of this work We therefore excluded these two methods from our evaluation and discussion.

#### S2.2 Proof of Theorem 1: The Unico 3D tensor estimator

*Proof.* Our goal is estimating $\{z_{ij}\}$, the realizations of $\{Z_{ij}\}$ given data $\{x_{ij}\}$ coming form $\{X_{ij}\}$ using $\mathrm{E}\left[Z_{ij}|\theta_j, w_i, X_{ij} = x_{ij}\right]$. We Consider a linear transformation $A_{ij} = Z_{ij} + B_{ij}X_{ij}$. If we construct it such that $A_{ij}, X_{ij}$ are non-linearly dependent then given $X_{ij} = x_{ij}$ we can express $Z_{ij}$ using the relation $A_{ij} - B_{ij}X_{ij}$ and calculate the conditional expectation as follows:

$$\begin{aligned}
\mathrm{E}\left[Z_{ij}|\theta_j, w_i, X_{ij} = x_{ij}\right] &= \mathrm{E}\left[A_{ij} - B_{ij}X_{ij}|\theta_j, w_i, X_{ij} = x_{ij}\right] && (6)\\
&= \mathrm{E}\left[A_{ij}|\theta_j, w_i, X_{ij} = x_{ij}\right] - \mathrm{E}\left[B_{ij}X_{ij}|\theta_j, w_i, X_{ij} = x_{ij}\right]\\
&= \mathrm{E}[A_{ij}|\theta_j, w_i] - B_{ij}x_{ij} = \mathrm{E}[Z_{ij}|\theta_j, w_i] + B_{ij}\left(\mathrm{E}[X_{ij}|\theta_j, w_i] - x_{ij}\right) && (7)
\end{aligned}$$

The above linear independence holds if we require that the cross-covariance vector between $A_{ij}$ and $X_{ij}$ is zero as follows:

$$\begin{aligned}
& \mathrm{Cov}[A_{ij}, X_{ij}] = (0, ..., 0) && (8)\\
\iff\ & \mathrm{Cov}[Z_{ij} + B_{ij}X_{ij}, X_{ij}] = (0, ..., 0) \iff \mathrm{Cov}[Z_{ij}, X_{ij}] + \mathrm{Cov}[B_{ij}X_{ij}, X_{ij}] = (0, ..., 0)\\
\iff\ & \mathrm{Cov}[Z_{ij}, X_{ij}] + B_{ij}\mathrm{V}[X_{ij}] = (0, ..., 0) \iff B_{ij} = -\mathrm{V}[X_{ij}]^{-1}\mathrm{Cov}[Z_{ij}, X_{ij}] && (9)
\end{aligned}$$

Noting that

$$\mathrm{Cov}[Z_{ijq}, X_{ij}] = \sum_{h=1}^{k} \mathrm{Cov}[Z_{ijq}, w_{ih}Z_{ijh}] = \sum_{h=1}^{k} w_{hi}\sigma_{jq,jh} \qquad (10)$$

and denoting $\Sigma_j \in \mathbb{R}^{k\times k}$ as the variance matrix of gene $j$ such that $(\Sigma_j)_{hq} = \sigma_{jh,jq}$, we thus get

$$B_{ij} = -\mathrm{V}[X_{ij}]^{-1}w_i^T\Sigma_j \qquad (11)$$

In summary, we get the following Unico estimator:

$$\hat{z}_{ij} = \mathrm{E}[Z_{ij}|\theta_j] + \mathrm{V}[X_{ij}]^{-1}\Sigma_j w_i (x_{ij} - \mathrm{E}[X_{ij}|\theta_j, w_i]) \tag{12}$$

$$\mathrm{E}[X_{ij}|\theta_j, w_i] = w_i^T \left(\mu_j + (c_i^{(1)})^T \gamma_j\right) + (c_i^{(2)})^T \beta_j \tag{13}$$

$$\mathrm{V}[X_{ij}|\theta_j, w_i] = \mathrm{Sum}\left((w_i w_i^T) \odot \Sigma_j\right) + \tau_j^2 \tag{14}$$

where $\gamma_j = (\gamma_{j1}, ..., \gamma_{jk}) \in \mathbb{R}^{p_1 \times k}$ is a matrix composed of the vectors $\{\gamma_{jh}\}$, the $\odot$ operator is the Hadamard product of two matrices, and the $\mathrm{Sum}(\cdot)$ operator is a summation across all entries of a matrix. $\square$

#### S2.3 Proof of Theorem 2: Improved capacity to reduce covariance bias

*Proof.* Consider a simple scenario where $\forall h : \mu_{jh} = 0, \sigma_{jh}^2 = 1$, we have no covariates, and $\tau_j = 0$ for some gene $j$. In this case, the TCA estimator becomes [6]:

$$\hat{z}_{ijh}^{\mathrm{TCA}} = \frac{w_{ih} x_{ij}}{\|w_i\|_2^2} \tag{15}$$

While TCA does not explicitly model cell-type covariance, we can use this estimator to calculate the covariance between the TCA-estimated values of two cell types $h \neq l$ (denote $\hat{\sigma}_{jh,jl}$) without observing an actual data point $x_{ij}$. For a given sample $i$ with $w_i$ we get

$$\mathrm{Cov}\left[\hat{z}_{ijh}^{\mathrm{TCA}}|X_{ij}, \hat{z}_{ijl}^{\mathrm{TCA}}|X_{ij}\right] = \frac{w_{ih} w_{il} \mathrm{V}[X_{ij}]}{\|w_i\|_2^4} = \frac{w_{ih} w_{il}}{\|w_i\|_2^2} \tag{16}$$

Now, while we treat the cell-type proportions $\{w_i\}$ as fixed for the samples in the data, they have some distribution in the population. Since cell-type proportions are bounded to the range $[0, 1]$ the mean of that distribution exists, hence, combining the information across samples we get

$$\hat{\sigma}_{jh,jl}^{\mathrm{TCA}} = \frac{1}{n} \sum_{i=1}^{n} \frac{w_{ih} w_{il}}{\|w_i\|_2^2} \xrightarrow{p} c_{hl} \tag{17}$$

where $c_{hl}$ is some constant, specific to cell types $h, l$ yet independent of $j$ and thus does not necessarily reflect the true correlation between the two cell types. In contrast, similarly considering the Unico estimator from Equations (12)-(14) under the same assumptions, we get:

$$\hat{\sigma}_{jh,jl}^{\mathrm{Unico}} = \frac{1}{n} \sum_{i=1}^{n} \frac{((\Sigma_j)_h^T w_i)((\Sigma_j)_l^T w_i)}{\mathrm{V}[X_{ij}]} \xrightarrow{p} f(\Sigma_j) \tag{18}$$

where $(\Sigma_j)_h$ is the $h$-th column of $\Sigma_j$ and $f(\Sigma_j)$ is some function of the variance matrix of gene $g$. It is easy to see that if $\Sigma_j = I_k$ then $f(\Sigma_j) = c_{hl}$ and Unico reduces to TCA. However, for genes with a non-trivial covariance matrix, the Unico estimator, in principle, has more capacity to alleviate the limitation of TCA in reflecting true covariances between cell types. $\square$

#### S2.4 Optimization of the Unico model

**Optimization setup.** We estimate the unknown parameters of the model by following concepts from the Generalized Method of Moments (GMM) [11]. The GMM framework allows us to learn the parameters of a model by solving equations (moment conditions) that match population moments (or, more generally, a function of population moments) with their corresponding data-derived sample moments. More concretely, let $f(\Theta, X_n) = f_1(\Theta, X_n), ..., f_d(\Theta, X_n)$ be $d$ moment conditions defined over parameters $\Theta$, where $X_n = (x_1, ..., x_n)$ are observations coming from the assumed model and $d > |\Theta|$. If the moment conditions satisfy $\forall 1 \leq l \leq d$ : $\mathrm{E}[f_l(\Theta, X_n)] = 0$ then an asymptotically consistent estimator of $\Theta$ is given by [11]:

$$\hat{\Theta} = \underset{\Theta}{\operatorname{argmin}}\, f(\Theta, X_n)^T \hat{U}_n f(\Theta, X_n) \tag{19}$$

where $\hat{U}_n$ is a positive definite (PD) weighting matrix. In the Unico model we treat $X_{ij}$ as a random variable with mean and variance that depend on gene-specific parameters ($\theta_j$), as well as on sample-specific cell-type proportions $w_i$. As a result, the variables $\{X_{ij}\}$, in general, have different distributions, however, these distributions are all parameterized by the same set $\theta_j$ of constant size. This leads us to construct a separate moment condition for each sample, which results in a multi-equation GMM problem with shared parameters across the Equation [12]. Since each moment condition is represented by a single sample the optimization of the Unico model eventually reduces to solving multiple weighted least squares problems.

Below, we provide a detailed derivation of Unico's optimization procedure. A summary of the optimization procedure is provided in Algorithm 1.

**Learning the means of the model.** We start by learning the means, which can be estimated independently from the variances and covariances. Specifically, let

$$\phi_j = (\mu_{j1}, ..., \mu_{jk}, \delta_{\mu_{j1}}, ..., \delta_{\mu_{jk}}, \gamma_{1j1}, ..., \gamma_{p_1jk})$$

be a vector of the means of gene $j$, for each sample $i$ in the data we define $f_i(\phi_j, x_{ij}) = x_{ij} - \mu_{X_{ij}}$, such that $\mu_{X_{ij}} \equiv \mathrm{E}[X_{ij}]$; clearly, $\mathrm{E}[f_i(\phi_j, x_{ij})] = 0$.

In theory, under a typical GMM estimation, setting the weights matrix $\hat{U}_n$ in Equation (19) to be the inverse of the empirical variance-covariance matrix of the moment conditions results in an asymptotically efficient estimator [11]. In our case, given that different moment conditions are based on independent individuals, this corresponds to setting $\hat{U}_n \in \mathbb{R}^{n\times n}$ as a diagonal matrix with values $\{\hat{u}_{ij} = (x_{ij} - \hat{\mu}_{X_{ij}})^{-2}\}_i$ on the diagonal; thus, reducing Equation (19) to the following weighted least squares problem for estimating $\phi_j$:

$$\hat{\phi}_j = \underset{\phi_j}{\operatorname{argmin}} \sum_{i=1}^{n} \hat{u}_{ij} \left(x_{ij} - \phi_j^T s_i\right)^2 \tag{20}$$

$$s_i = (w_{i1}, ..., w_{ik}, w_{i1}c_{i1}^{(1)}, ..., w_{ik}c_{ip_1}^{(1)}, ..., c_{i1}^{(1)}, ..., c_{ip_1}^{(1)}, c_{i2}^{(2)}, ..., c_{ip_2}^{(2)}) \tag{21}$$

Indeed, we know from large sample theory of weighted least squares problems that this is an asymptotically consistent and efficient estimator. However, in practice, we choose to forfeit asymptotic efficiency for practical considerations in finite data. Specifically, we wish to avoid very high values of $\hat{u}_{ij}$, which can greatly affect the solution by relying on just a small number of data points with extreme weights. In practice, we address this by modifying the definition of $\hat{u}_{ij}$ to allow only down-weighting of samples:

$$\hat{u}_{ij} = \min\left(1, (x_{ij} - \hat{\mu}_{X_{ij}})^{-2}\right) \tag{22}$$

Notably, Equation (22) relies on an estimate of $\mu_{X_{ij}}$, which requires estimating the parameters $\phi_j$. However, these are exactly the parameters that we wish to estimate in Equation (20). We resolve this by employing an alternating optimization: we first solve Equation (22) while setting $\forall i : \hat{u}_{ij} = 1$, and then, given initial estimates of $\phi_j$, we follow Equation (22) for constructing the weights for subsequent optimization of Equation (22). This procedure can be followed by more iterations until convergence, much like in a standard feasible generalized least squares.

Solving Equation (20) requires us to learn a total of $k(1+p_1)+p_2$ parameters for each gene. This is a constant number of parameters compared with the number of data points used for learning these parameters ($n$). The optimization of each gene is independent, which makes the problem feasible in principle, however, in practice, any application of the above optimization should be mindful of the balance between the sample size and this number of parameters.

**Learning the variances and covariances of the model.** Once the algorithm converges to a set of means ($\hat{\phi}_j$), we move on to the estimation of the variances and covariances of gene $j$. Let $\psi_j = \text{diag}(\Sigma_j)$ and let $\xi_j = \text{tri}(\Sigma_j)$ be a vector of the elements in the lower diagonal entries of the matrix $\Sigma_j$ (excluding the diagonal elements). We follow the same approach we take for estimating the means, which yields the following optimization problem:

$$\hat{\psi}_j, \hat{\xi}_j, \hat{\tau}_j^2 = \underset{\psi_j, \tau_j^2}{\operatorname{argmin}} \sum_{i=1}^{n} \hat{v}_{ij} \left( (x_{ij} - \hat{\mu}_{X_{ij}})^2 - \psi_j^T s_i' - 2\xi_j^T s_i'' - \tau_j^2 \right)^2 \tag{23}$$

$$s_i' = w_i \odot w_i, s_i'' = \operatorname{tri}(w_i w_i^T) \tag{24}$$

$$\hat{v}_{ij} = \min\left(1, \left((x_{ij} - \hat{\mu}_{X_{ij}})^2 - \hat{\sigma}_{X_{ij}}^2\right)^{-2}\right), \quad \sigma_{X_{ij}}^2 \equiv \mathrm{V}[X_{ij}] \tag{25}$$

In order to direct the inference towards a good solution we further wish to constrain $\hat{\psi}_j, \hat{\xi}_j$ to form an estimate of $\Sigma_j$ that is positive definite (PD). We, therefore, reformulate the above optimization problem as follows. Consider the Cholesky decomposition $\Sigma_j = L_j L_j^T$ where $L_j$ is a lower triangular matrix with positive values on its diagonal. Instead of directly estimating the elements of $\Sigma_j$ by solving Equation (23)-(25), we guarantee that $\hat{\Sigma}_j$ is PD by estimating the non-zero elements of $L_j$. Specifically, we express $\Sigma_j$ in the optimization problem through $L_j$, which results in the following optimization problem:

$$\hat{L}_j, \hat{\tau_j}^2 = \underset{L_j, \tau_j^2}{\operatorname{argmin}} \sum_{i=1}^{n} \hat{v}_{ij} \left( (x_{ij} - \hat{\mu}_{X_{ij}})^2 - d_j^T s_i' - 2 l_j^T s_i'' - \tau_j^2 \right)^2 \tag{26}$$

$$\text{s.t.}, \forall h \in \{1, ..., k\} : L_{hh} > 0 \tag{27}$$

where

$$d_j = \operatorname{diag}(L_j L_j^T), l_j = \operatorname{tri}(L_j L_j^T) \tag{28}$$

$$s_i' = w_i \odot w_i, s_i'' = \operatorname{tri}(w_i w_i^T) \tag{29}$$

$$\hat{v}_{ij} = \min\left(1, \left((x_{ij} - \hat{\mu}_{X_{ij}})^2 - \hat{\sigma}_{X_{ij}}^2\right)^{-2}\right), \quad \sigma_{X_{ij}}^2 \equiv \mathrm{V}[X_{ij}] \tag{30}$$

Finally, given $\hat{L}_j$, we set $\hat{\Sigma}_j = \hat{L}_j \hat{L}_j^T$. The above problem is non-convex, however, it requires learning small independent problems of $\binom{k}{2} + k + 1$ parameters for each gene (where $k$ is a small constant; e.g., 5 cell types). Similarly to the estimation of the means, we apply an alternating optimization procedure to overcome the need for an estimate $\hat{\sigma}_{X_{ij}}^2$ in Equation (30).

**Algorithm 1** Optimization procedure for **Unico**. Notations are based on subsection S2.4

**for** each feature $j$ present in genomic data $X$ **do**
    $\forall i \quad \hat{u}_{ij} \leftarrow 1$ ▷ Initialize weights
    **while** `not converged` **do** ▷ Estimate the means
        $\hat{\phi}_j \leftarrow \text{argmin}_{\phi_j} \sum_{i=1}^{n} \hat{u}_{ij} \left( x_{ij} - \phi_j^T s_i \right)^2$
        **for** each sample $i$ **do**
            $\hat{u}_{ij} \leftarrow \min \left( 1, (x_{ij} - \hat{\mu}_{X_{ij}})^{-2} \right)$
        **end for**
    **end while**
    $\forall i \quad \hat{v}_{ij} \leftarrow 1$ ▷ Initialize weights
    **while** `not converged` **do** ▷ Estimate variances and covariances
        $\hat{L}_j, \hat{\tau_j}^2 \leftarrow \text{argmin}_{L_j, \tau_j^2} \sum_{i=1}^{n} \hat{v}_{ij} \left( (x_{ij} - \hat{\mu}_{X_{ij}})^2 - d_j^T s_i' - 2 l_j^T s_i'' - \tau_j^2 \right)^2$
        $\hat{\Sigma}_j \leftarrow \hat{L}_j \hat{L}_j^T$
        **for** each sample $i$ **do**
            $\hat{v}_{ij} \leftarrow \min \left( 1, \left( (x_{ij} - \hat{\mu}_{X_{ij}})^2 - \hat{\sigma}_{X_{ij}}^2 \right)^{-2} \right)$
        **end for**
    **end while**
**end for**

#### S2.5 Generating pseudo-bulk mixtures from scRNAseq profiles

There are currently no large publicly available bulk datasets with matching cell-type level data for the same group of individuals. We therefore simulated pseudo-bulk PBMC expression profiles using single-cell PBMCs from the Stephenson et al. study [4] and pseudo-bulk lung expression profiles using data from the Human Lung Cell Atlas (HLCA) [13]. In order to account for sequencing depth, gene counts were converted to counts-per-million (CPM) in every cell before averaging all cells of type $h$ from sample $i$ to generate a cell-type-specific pseudo-bulk expression pattern.

We estimated per-sample cell-type proportions for both single-cell datasets by calculating cell counts per individual sample for all modeled cell types ($k \in \{5, 7\}$ for PBMC and $k \in \{4, 6\}$ for HCLA) and normalizing them to sum up to 1 per sample. Average proportions were 31.8%, 19.5%, 17.7%, 16.7%, 14.2% for the top five main cell types in PBMC: CD4 T cells, NK cells,

CD8 T cells, monocytes and B cells, respectively (k=5 scenario). Monocytes can be further stratified to 14.7% CD14 monocytes and 2.0% CD16 monocytes, and B cells can be further stratified to 13.4% canonical B cells and 0.8% plasma cells (k=7 scenario). In the lung data, we only considered samples collected from parenchyma tissues, which demonstrated average proportions of 62.5%, 23.8%, 8.4%, and 5.4% for the top four main cell types: immune cells, epithelial cells, endothelial cells, and stromal cells, respectively (k=4 scenario). Immune cells can be further stratified to 43.5% myeloids and 17.4% lymphoid cells, and epithelial cells can be further stratified to 18.8% alveolar epithelium and 5.6% airway epithelium cells (k=6 scenario).

We generated every pseudo-bulk sample by mixing cell-type proportions from one sample (randomly drawn with replacement) and all cell-type profiles of a single sample (randomly drawn with replacement). In addition, we drew a noisy version of the cell-type proportion estimates from the following Dirichlet distribution: $\mathrm{Dir}(\alpha w_i)$, where $w_i$ denotes the ground truth proportion of the sample and $\alpha \in \{100, 50, 25, 10, 5, 2.5\}$ controls the noise level. Large $\alpha$ forces the distribution to be concentrated around the true proportion while smaller ones allow more derivation. To avoid sampling extremely lowly expressed genes, we restricted our sampling space to the top 10,000 most expressed genes, evaluated based on the gene-specific sum of average expression across cell types. In cases where genes demonstrated too low expression in a certain cell type (mean or variance less than $10^{-4}$ after excluding outliers that are 2 standard deviations away from the mean), we added a small non-negative Gaussian noise $\mathcal{HN}(0, 10^{-4})$. This was done to improve numerical stability and to ensure the gene-level correlation matrix remains of full rank when calculating cell-type covariance entropy. Per simulated dataset, we carried out the above-mentioned sampling strategy independently for 600 genes, and for varying numbers of sample sizes (100, 250, or 500 samples). Per experiment, we repeated the mixture sampling procedure 20 times.

For the purpose of evaluation, both the pseudo-bulk mixtures and cell-type specific expression profiles were scaled by the standard deviation of each gene, which we calculated from the pseudo-bulk data so that the variance across different genes is roughly comparable. Omitting this standardization step would induce spuriously high correlation scores for algorithms that simply estimate the relative scale of the parameters correctly.

Finally, for every cell type, we excluded genes with associated cell-type expression profiles that demonstrated (after scaling) mean or variance $\leq 0.1$ from correlation calculation and multiple linear regression analysis with pseudo-bulk profile. This exclusion alleviated the risk of evaluating deconvolution methods against merely artificially added noise as described earlier or extremely noisy counts. Of note, in general, CIBERSORTx estimates cell-type proportions from the bulk input. Here, however, we directly provided it with the ground truth cell-type proportion of the mixtures, as provided to all other methods we benchmarked.

#### S2.6 Cell-type level differential methylation analysis

The Unico model naturally allows statistical testing for cell-type level associations by incorporating a phenotype of interest as a cell-type level covariate $\{c_i^{(1)}\}$ and testing whether its effect is non-zero (i.e., whether $\gamma_{jh} \neq 0$ for a given feature $j$ and cell-type $h$). This does not require us to explicitly estimate the underlying 3D tensor of cell-type levels. Instead, we can estimate effects directly following the model in Equations (1)-(5). As we describe below, we can take either a parametric approach or a non-parametric approach for such statistical testing.

##### S2.6.1 Statistical testing with Unico under a parametric assumption

In our analysis of differential methylation (DM), we assume that cell-type methylation levels are normally distributed, which amounts to the following assumption under the Unico model:

$$Z_{ij} \sim \mathcal{N}\left(\mu_j + (c_i^{(1)})^T \gamma_{jh}, \sigma_{jh}^2\right) \tag{31}$$

$$X_{ij} \sim \mathcal{N}\left(w_i^T(\mu_j + (c_i^{(1)})^T \gamma_{jh}) + (c_i^{(2)})^T \beta_j, \mathrm{Sum}\left((w_i w_i^T) \odot \Sigma_j\right) + \tau_j^2\right) \tag{32}$$

For a given CpG $j$ under test, Equation (32) corresponds to a heteroskedastic regression problem with $\{x_{ij}\}_i$ as the dependent variable and $\{w_i\}, \{w_i c_i^{(1)}\}, \{c_i^{(2)}\}$ as the independent variables. This view allows us to perform statistical testing by solving a generalized least squares problem using a standard linear regression framework. Concretely, we scale every methylation sample $i$ by the inverse of its estimated standard deviation:

$$q_{ij} := \left(\mathrm{Sum}\left((w_i w_i^T) \odot \hat{\Sigma}_j\right) + \hat{\tau}_j^2\right)^{-0.5} \tag{33}$$

where $\hat{\Sigma}_j, \hat{\tau}_j^2$ correspond to the variance and covariance parameters estimated under the non-parametric model optimization of Unico. More specifically, we scale both the dependent and independent variables of sample $i$ by $q_{ij}$ and fit a standard linear regression model. Statistical testing then becomes straightforward: marginal cell-type level effects can be evaluated using a standard t statistic, and tissue-level effects can be evaluated using a partial F statistic that quantifies the joint effect across all cell types.

##### S2.6.2 Distribution-free statistical testing with Unico

Given the estimated model parameters (means, variances, and covariances), we can derive asymptotic p-values for cell-type-specific effect sizes. Let $S$ be a design matrix formed by stacking $s_1, s_2, ..., s_n$ following Equation (21) as the rows, and let $Q_j = \text{diag}(q_{1j}^2, ..., q_{nj}^2)$ be a diagonal weighting matrix, where $q_{ij}$ follows Equation (33), we solve:

$$\hat{\phi_j}^{\text{asym}} = \underset{\phi_j}{\text{argmin}}(x_j - S\phi_j)^T Q_j (x_j - S\phi_j) \tag{34}$$

where $x_j = (x_{1j}, ..., x_{nj})$. We get a weighted least squares (WLS) problem, which is characterized by the following analytical solution and asymptotic distribution of the estimator $\hat{\phi_j}^{\text{asym}}$:

$$\hat{\phi_j}^{\text{asym}} = (S^T Q_j S)^{-1} S^T Q_j x_j \tag{35}$$

$$\hat{\phi_j}^{\text{asym}} \xrightarrow{d} N(\phi_j, (S^T Q_j S)^{-1} S^T Q_j \text{V}[f(\phi_j, X_{ij})] Q_j S (S^T Q_j S)^{-1}) \tag{36}$$

where $f(\phi_j, X_{ij}) = x_{ij} - \mu_{X_{ij}}$ and its empirical variance is $\text{V}[f(\phi_j, X_{ij})] = (x_{ij} - \mu_{X_{ij}})^2$. This can be written more compactly as:

$$\hat{\phi_j}^{\text{asym}} \xrightarrow{d} N(\phi_j, (S^T Q_j S)^{-1} S^T Q_j^* S (S^T Q_j S)^{-1}) \tag{37}$$

$$(Q_j^*)_{ii} = (Q_j)_{ii}^2 (x_{ij} - \mu_{X_{ij}})^2 \tag{38}$$

where $(Q_j)_{ii}$ denotes the $i$-th element on the diagonal of $Q_j$. Now, following Slutsky's Theorem, we get:

$$\frac{(\hat{\phi_j}^{\text{asym}})_p}{\text{SE}(\hat{\phi_j}^{\text{asym}})_{pp}} \xrightarrow{d} N(0, 1) \tag{39}$$

$$\text{SE}(\hat{\phi_j}^{\text{asym}})_{pp} = \sqrt{[(S^T Q_j S)^{-1} S^T Q_j^* S (S^T Q_j S)^{-1})]_{pp}} \tag{40}$$

where $(\hat{\phi_j}^{\text{asym}})_p$ denotes the $p$-th entry of $\hat{\phi_j}^{\text{asym}}$. Given Equation (39), statistical testing becomes straightforward using the cumulative distribution function of the standard normal distribution. Since the moment condition $f(\hat{\phi_j}, X_{ij})$ is defined based on a single data point, we excluded samples with very small empirical variance $V[f(\hat{\phi_j}, X_{ij})]$; otherwise, those could dominate the solution. Concretely, we excluded 5% of the samples with the lowest empirical variance.

##### S2.6.3 Statistical testing using competing deconvolution methods

Testing for DM with sex and age should consider the effect of these demographics on methylation levels and not vice versa [7]. Assuming this model directionality is natural within the Unico model, which can evaluate the effect of sex and age as cell-type level covariates in (3). Similarly, we applied TCA, CellDMC, and our baseline model to evaluate the effects of sex and age on cell-type level methylation. bMIND, on the other hand, required a different treatment. The bMIND implementation includes two approaches for association testing (both of which only support binary phenotypes; i.e., only sex in our case). The first approach models the appropriate direction of effect, as indicated above, however, it relies on MCMC sampling, which renders it infeasible for epigenome-wide association studies; particularly, in our case, execution time is expected to be over 24 hours for a single CpG assuming 30 threads in order to pass a Bonferroni-corrected threshold. We thus opted for the alternative, computationally feasible approach, which the authors of bMIND also recommended. Specifically, we performed testing directly on the estimated tensors, which implicitly makes a non-sensible assumption that methylation affects sex. To that end, cell-type level effects and corresponding p-values were derived by fitting a logistic regression model with cell-type level profiles of the estimated tensor and covariates as the independent variables and sex as the dependent variable.

In addition to the evaluation of cell-type-level DM, we further tested for tissue-level DM. Similarly to Unico, TCA allows calculating such tissue-level p-values [6]. In contrast, CellDMC only evaluates and reports summary statistics for marginal cell-type level tests [14]. We, therefore, implemented a tissue-level test for CellDMC by learning a restricted linear regression model without the cell-type interaction terms and performing a partial F-test based on the original unrestricted model. For bMIND, p-values were derived for the tissue-level tests from a multivariate analysis of covariance (MANCOVA) with sex as the independent variable and cell-type level profiles as dependent variables with $\{c_i^{(1)}\}$ and $\{c_i^{(2)}\}$ covariates adjusted. Finally, when evaluating tissue-level DM based on a straightforward analysis of the bulk data, we performed a standard linear regression analysis directly on the bulk mixture with bulk data as the dependent variable and cell-type proportions, as well as all covariates, as independent variables. P-values for tissue-level effects were derived using a standard t statistic.
